# Mind the gap: assessing the co-occurrying impacts of intensification and land abandonment on bird diversity in a farmland mosaic of Northeastern Portugal using historical data (1980–2000)

**DOI:** 10.1101/2023.11.10.566211

**Authors:** Tiago Múrias, Sérgio B. Ribeiro, Angela Lomba, Nuno Ferrand De Almeida

## Abstract

Agriculture intensification and land abandonment are the main threats to bird communities in farmlands. Although they represent the opposite endpoints of a gradient of land-use change, they are usually examined separately. In this study, we assessed the effects of the occurrence of both drivers in space and in time, on bird assemblages. The study was performed in a Mediterranean vineyard-dominated farmland mosaic located in North-Eastern Portugal, where land abandonment and (mechanized) intensification coexisted for twenty years. The aims of this study were (1) to illustrate and describe the temporal patterns of land-use change; (2) to investigate whether these changes were associated with changes in the species composition structure of the local bird communities;, (3) to assess which species/ecological traits were more responsive to land-use changes. We compared the occurrence of birds before (1979-84) and after (2002-03) the triggering of the joint land-use change process. Results showed the prevalence of two potentially interacting directions of land-use change: land intensification and land abandonment. While the composition and ecological structure of the bird communities remained stable during the period analyzed, trait and species-specific analysis showed significant correlations with the gradients of change. This suggests that the local bird communities were able to cope with the synergic dynamics of the land-use change, possibly due to a combination of a resilient pre-change bird assemblage and the buffering effect of the competing drivers. The pros and cons of the approach adopted and the utility of past datasets as baselines to assess and monitor the impacts of environmental change on farmland bird diversity are discussed.

## 1. Introduction

The ongoing decline of farmland in Europe is well documented (Donald et al., 2001, 2006; Vorisek et al., 2010; Sanderson et al., 2013) and is a source of increasing concern due to the value of these human-shaped ecosystems to farmland birds, some of conservation concern (Birdlife International, 2004; Parachinni et al, 2008). This decline has been related to the effect of two contrasting land-use trends: agricultural intensification (here defined as the mechanization of the productive proccess) and land abandonment (i.e. the cessation of agricultural management) (Stoate et al., 2001, Russo, 2006; Sanderson et al., 2013).

Whilst the effects of intensification on farmland birds are generally negative (Donald et al., 2001, 2006; Reif et al., 2008; Sanderson et al., 2013; Jeliazkov et al., 2016), the effects of land abandonment are more fuzzy, as different species are impacted differently (Russo, 2006; Herrando et al., 2014; Regos et al., 2016), according to species-specific ecological traits (Parodi et al., 2001; Sirami et al., 2007; Brambilla et al., 2010) or biogeographical origin (Suárez-Seoane et al. 2002). In fact, the loss of farmland species due to abandonment can be numerically counterbalanced by scrubland or forest species (Russo, 2006; Keenleyside and Tucker, 2010; Herrando et al., 2014), maintaining the overall species richness despite underlying species turnover. Nevertheless, both agricultural intensification and land abandonment ultimately drive the degradation of suitable habitats for farmland specialists, reshaping species composition and promoting the biotic homogenization of avian communities, where the increasing dominance of a few species leads to a general decrease of species diversity (Flynn et al., 2009; Clavel et al., 2011; Guerrero et al., 2011).

Closely linked to the Common Agriculture Policy (CAP) (Donald et al., 2002), agriculture intensification has been observed in the most productive lowland areas of North Western Europe (Donald et al., 2001, 2006; Stoate et al., 2001). Similar trends have been described in several countries of Central and Eastern Europe after their accession to the European Union (EU) (Reif et al., 2008; Sanderson et al., 2013; Reif and Vermouzek, 2018). Conversely, land abandonment typically prevails in less productive areas, generally in remote mountain regions (some designated as Less Favoured Areas; Merckx & Pereira, 2015), of Southern and SE Europe, where poor soils and harsh climate prevail (Stoate et al., 2009; Keenleyside and Tucker, 2010; Zakkak et al., 2015). Due to this (apparent) spatial segregation, these drivers have been examined separately in the European context. However, both land-use trends are extremes of a gradient of agricultural management whose spatial and temporal dynamics depend on the social-ecological context, and are reflected in specific landscape patterns linked to the evolution of the socio-economic pressures and thus likely to co-occur in space and in time (eg., Katayama et al., 2015).

Assessing the response of farmland birds to the impacts of land-use change is challenging, due to the complexity and long-term nature of the process. Typically, studies dealing with this research topic in Europe use regression models to relate the birds abundances (or frequency of occurrence) to relevant environmental drivers (Reif et al., 2008; Nikolov, 2010; Herrando et al., 2014). Overall, such studies build on data on birds’ abundance collected in the context of long-term censuses at national or European scales, like the Pan-European Common Bird Monitoring Scheme (PECBMS, 2019). However, required long-term data is scarce and restricted to a few European countries (Siriwardena et al., 1998; Wretenberg et al., 2007; Jiguet et al., 2012), or to specific “time windows” (Scozzafava and Santis, 2006; Sirami et al., 2008; Fonderflick et al., 2010). A common practice to overcome the absence of such data, is the usage of a “space-for-time” method, in which spatial plots in different phases of the ecological succession are surveyed at the same time and used as temporal surrogates (Moreira et al., 2001; Scorzaffava and Sanctis, 2006). Extrapolating from such data, however, must be made with care due to potential problems of site representativeness and/or non-linear bird-area relationships (Moreira et al., 2001, Bonthoux et al., 2013; De Palma et al., 2018). An increasing popular, and cost-effective alternative is to collate data from bird atlases published in the past decades, whose utility for conservation purposes, despite known limitations, has been demonstrated (Gibbons et al, 2007; Dunn and Weston, 2008; Robertson et al., 2010).

In Portugal, data on the relationships between birds and habitat change in farmland landscapes is relatively recent compared to other countries (Delgado and Moreira, 2000; Moreira et al., 2005; Reino et al; 2010). In fact, analyses of temporal trends don’t include the period of the pre-1993 CAP reform (Ribeiro et al. 2014; Santana et al., 2014, 2017). Moreover, such studies target almost exclusively High Nature Value (HNV) cereal pseudo-steppes and cork woodlands of southern Portugal, whereas the problem of intensification and/or land abandonment in the mountain farmland mosaics above the Tagus River (northern Portugal) have seldom been addressed (Moreira et al., 2001; Guilherme, 2013). We aim to fill this knowledge gap by using information from an unpublished local bird atlas performed in 1979-84 (Ferrand de Almeida, unpublished) and repeated in early 2000’s (T. Múrias, unpublished data), to assess the potential impacts of land-use change on a typical mountain farmland bird community of northeastern Portugal.

The study area is located in the Upper Douro Wine Demarcated Region (*Região Demarcada do Douro,* RDD), the most important wine-producing region of Portugal (Andresen et al., 2005). The RDD was characterized, in large part and for a long time, by the predominance of small farms using non-mechanized methods of production (Barreto, 1992). Since the 1960’s, and following a national pattern, a continuous drainage of manpower and increasing economic unprofitability led to the abandonment of production in many of these small farms. However, between 1980 and 2000, due to EU’s incentives for the wine sector, many of the previously abandoned farms were reactivated or revitalized using more profitable (mechanized) agricultural management practices (Barreto, 1992). These dynamics promoted the co-occurrence of both intensification and land abandonment in space and in time, providing an excellent opportunity to explore the potential of their combined effects on the local bird communities in the study area. In this context, we postulate that the co-occurrence of intensification and land abandonment should exacerbate their respective individual effects which ultimately would be reflected as a significant loss of farmland specialists (due to intensification) and a small to moderate increase in forest generalists (due to land abandonment), with the net effect being a general impoverishment of the bird community composition.

Thus, the aims of this study were (1) to characterize the spatio-temporal trajectories of land-use change in the study area between 1980 and 2000 in order to detect potential signals of both drivers, and how did they operate in the region; (2) to examine potential shifts in the composition and structure of local bird assemblages in response to landscape change, namely, evaluating whether landscape shifts driven by agricultural restructuring and abandonment led to biotic homogenization, mediated by expanding generalist species (community convergence), or to local community differentiation (community divergence), and (3) to determine which functional ecological traits (and species) explained the responses of local assemblages to land-use dynamics between 1980 and 2000. Results are discussed within the broader context of the utility of historical atlas-type data (i.e., presence/absence information) for establishing ecological baselines to assess and monitor the impacts of environmental change on farmland bird diversity.

## 2. Methods

### 2.1 Study area

The study area (41°15’ N, 7°24’ W, Figure 1A, rigth) is a 5000 ha Mediterranean mountainous farmland mosaic with hot summers and cold but wet winters (Ribeiro, 2000), located in the mouth of river Tua (northeast Portugal), within the limits of the Douro Wine Demarcated Region (“Região Demarcada do Douro”, RDD) (Figure 1A, left). Altitude ranges from 100 m near the riverbed to 600-700 m in the highest *plateaus*. The landscape is dominated by vineyards (*Vitis vinifera*) arranged in terraces bordered either by narrow schist walls (traditional vineyards), or larger earth walls (new mechanized or semi-mechanized vineyards) (Andresen et al., 2005). Other relevant cultures include traditionally managed olive-orchards, orange-groves and gardens. A variety of shrubs (*Juniperus oxycedrus* L.*, Lavandula stoechas* L., *Genista* spp., *Ulex* spp. And *Erica* spp.) and native oak-type woodlands (*Quercus suber* L. *and Q. rotundifolia* Lam.) occur in the slopes leading to river Tua and its associated tributary streams, while the hilltops and middle slopes are generally occupied by pinewoods (*Pinus pinaster* Aiton). A highland wet meadow is also present in the eastern margin of the river (Figure 1B).

**Figure 1.**
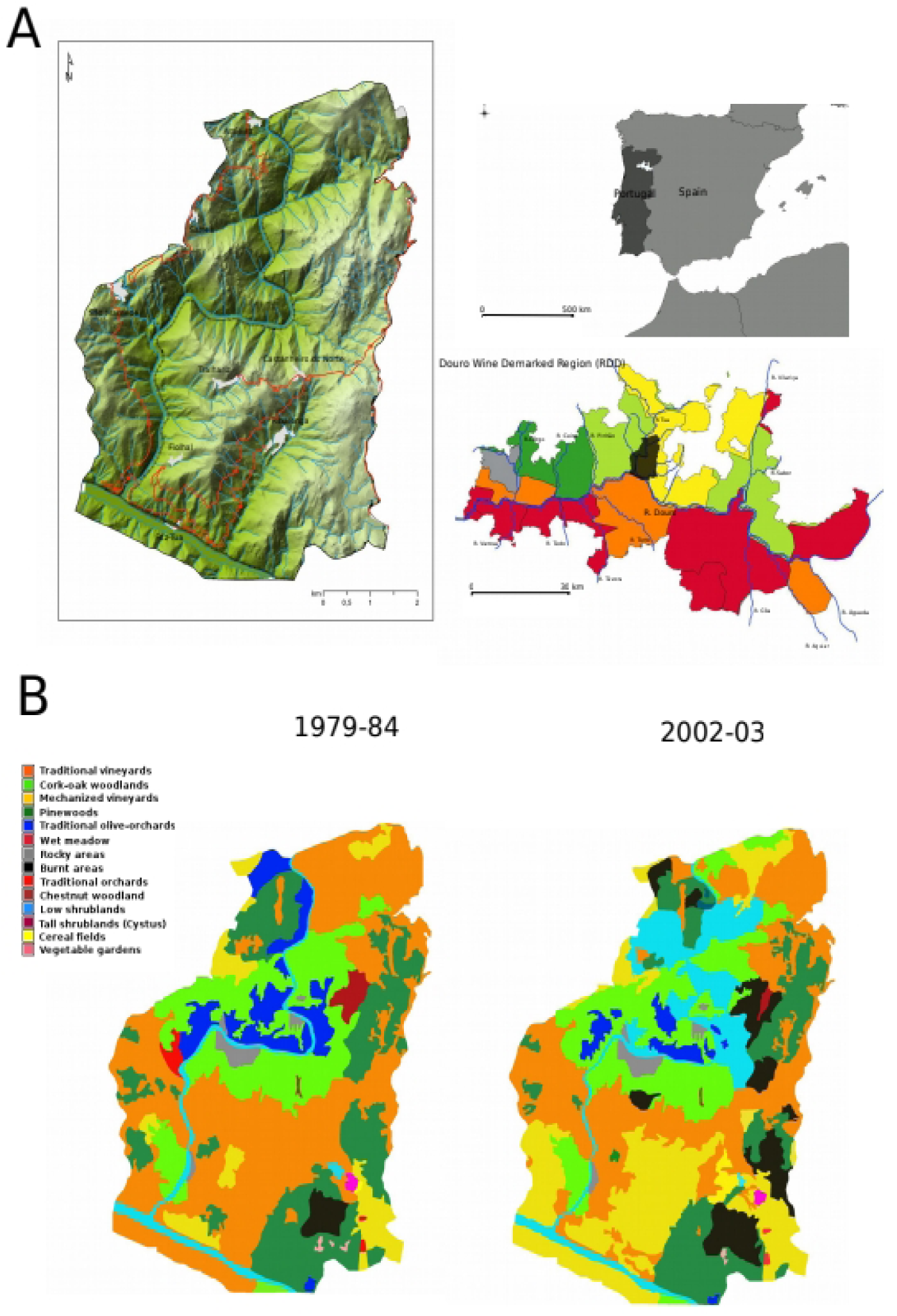
(A) Location of (1) the ‘Douro Wine Demarked Region’ ***(Região Demarcada do Douro,*** RDD) in Portugal, (2) of the study area in the RDD and (3) detailed Digital Terrain Model of the study area. The colours in Figure A.2 represent the gradient from heavy losses to medium gains in the total number of farms per ha of Usable Agricultural Area (farms/UAA) in the RDD counties between 1989 and 1999. Large decreases in the total number of farms per area (between 35% and 20% loss) are depicted in red; moderate decreases (19% to 10%) in orange; small decreases (9% to 1%) in yellow; no decreases or small gains (0% to 10% of gains) in ligth green; moderate gains (11% to 25%) in dark green; no data is available to the period concerned, in grey. (B) Spatial changes in land use in the study area between 1979-84 and 2002-03.

The human population in the study area is scarce, withthe majority of its 2000 inhabitants concentrated in eight small villages (INE, 2002; Figure 1A, left). Rural depopulation has been observed since the early 1960’s (PORDATA, 2018).

#### 2.2.1 Original data dataset

In the context of an Atlas of Terrestrial Vertebrates carried out between 1979 and 1984 (hereafter designated as “1980”), the study area was surveyed using the point census method described by Blondel (1969), adapted for presence/absence data (Ferrand de Almeida, unpublished). For this purpose, a grid of 231 500x500 m squares, covering the whole study area was established. From this total, a sample of 139 squares (60.2% of the total number), stratified by landscape class, was surveyed (see Figure S1A, Supplemantary Material; see Table 1 for a description of the landscape classes used).

**Table 1.** Approximate area of each land-use type in 1980 and 2000, net area variation (in hectares), percentage change between the two periods, and loadings along the land-use change gradients (Principal Component Analysis). Shrublands were dominated mostly by Cistus spp. and Genista spp. Percentage change is expressed relative to the initial area of each specific land-use class in 1980 (“by land-use type”) and relative to the total area of the major landscape unit (i.e., “agricultural” or “forest”) to which it belonged in 1980 (“by major landscape unit”). Only PCA axes with eigenvalues >1 are shown. Loadings of |0.50| and above (rounded up) are depicted in bold. The proportion of variance accounted for is calculated relative to the total variance across all nine original PCA axes, but only the results for the first four are shown.

| Land-use characterization (1980 - 2000) |  |  |  |  |  |  | Gradients of land-use change (PCA) |  |  |  |
| --- | --- | --- | --- | --- | --- | --- | --- | --- | --- | --- |
| *Land-use types* | *Code* | *Area (ha)* | | $\Delta$ (ha) | $\Delta$ (%) | | PC1 | PC2 | PC3 | PC4 |
|  |  | (1980) | (2000) |  | By land use type | By major landscape unit |  |  |  |  |
| **Agricultural landscapes** |  | **2797** | **2590** | **-207.0** | - | -7.4 |  |  |  |  |
| Traditional vineyards<sup>1</sup> | OV | 2092 | 1364 | -728 | -34.8 | -26.0 | **0.779** | 0.193 | **0.500** | 0.060 |
| Intensive vineyards<sup>2</sup> | Vin | 320 | 1101 | +781 | >100.0 | +27.9 | **-0.824** | -0.135 | -0.432 | -0.163 |
| Olive-orchards | Oli | 345 | 117 | -228 | -66.0 | -8.2 | **-0.528** | **0.57** | 0.203 | 0.375 |
| Other cultures<sup>3</sup> | Cul | 40 | 8 | -32 | -80.0 | -1.1 | -0.106 | -0.059 | **-0.502** | 0.168 |
| **Forest landscapes<sup>4</sup>** |  | **2262** | **2374** | **+112.0** | - | +0.5 |  |  |  |  |
| Pinewoodland | Pin | 1101 | 658 | -443 | -40.2 | -15.6 | -0.085 | **-0.664** | 0.243 | **0.658** |
| Oakwoodland | Cor | 868 | 833 | -35 | -4.0 | -1.5 | -0.312 | 0.063 | **0.626** | -0.339 |
| Meadow | Mea | 51 | 9 | -42 | -82.4 | -1.8 | -0.205 | -0.430 | 0.295 | **-0.540** |
| Shrublands | Shr | 173 | 533 | +360 | >100.0 | +15.2 | 0.628 | -0.527 | -0.499 | -0.151 |
| Burnt areas | Bur | 69 | 341 | +273 | +100.0 | +11.5 | 0.290 | 0.820 | -0.383 | -0.112 |
| Eigenvalues |  |  | - |  |  |  | 2.23 | 1.97 | 1.66 | 1.07 |
| Variance per axis ( % ) |  |  | - |  |  |  | 24.81 | 21.87 | 18.48 | 11.92 |
| Cumulative variance ( % ) |  |  | - |  |  |  | 24.81 | 46.69 | 65.16 | 77.08 |

<sup>1</sup> Traditional manned vineyards with scattered olive-trees in old terraces <sup>2</sup> Semi-intensive and intensive vineyards, mostly or fully mechanized <sup>3</sup> Orchards and gardens, mainly located around the villages and farms <sup>4</sup> *Sensu latu*, thatis including shrublands and other non-agricultural areas

Each selected square was visited at least once per season (spring, summer, winter and autumn) and year during the study period (range of 4 to 24 visits per square/period). Each sampling occasion lasted from 4-6 days (Figure S2C, Supplementary Material). The results for each species distribution in the sampled squares were then extrapolated to the entire study area, based on the species habitat use, ecological and biological requirements and the author’s knowledge of the local ecosystems (Ferrand de Almeida, unpublished). The final species maps thus included both recorded and potential presences (Figure S2A-B, Supplementary Material). They were complemented by a short description of the species primary habitat(s) and (local) phenology and an abundance estimate (following a semi-quantitative scale) in the study area. The habitat characterization mentioned above was used exclusively as a species ecological trait in the community-level and fourth-corner (4cp) analyses (see below). All spatial landscape metrics were strictly derived from the land cover units.

A land cover map for the study area with the main land-use classes, based on a previous photo-interpretation of 1986 aerial photos (spatial resolution 1:10.000), was also included (Figure S2D, Supplementary Material).

#### 2.2.2 Field surveys (2002-03)

In 2002/2003 (hereafter, designated as ‘2000’) a new survey was conducted using the same pool of squares sampled in 1979-84. Due to several logistic constraints (eg. lack of access or security issues) the number of plots visited in the second survey declined to 112 (80.6% of the area sampled in 1979-84 and 48.5% of the total number of squares), which was the final sample size used for the comparative analysis presented. However, this reduced sample still represented all landscape classes present.

Each square was visited at least once (but usually two times) over the study period (winter: November-February, spring: March-June and summer: July). The autumn was excluded due to the low detection probability in this period. This led to the removal of three strict autumn migrant species present in the 1979-84: *Muscicapa striata, Ficedula hypoleuca and Saxicola rubetra*. In each visit, all species seen or heared were recorded. The observers remained in the sampling point until no new species were detected, for a minimum period of 10 minutes and a maximum of 40 minutes. Coupled with the fact that most observations were auditive, this method contributed to maximize the species detection, ensuring a “saturation” of species in each square and sampling occasion. Species not detected during these standardized survey periods were considered absent for the purpose of the analyses (i.e., non-detections), as in the previous survey.

To evaluate whether this concentrated two-year sampling protocol provided robust coverage compared to the earlier extended (six-year) survey period, we formally tested sampling effort standardization using the neighborhood framework proposed by Hill (2012). Based on the complete baseline matrix from the first campaign (n=139 squares), a dedicated R script implementing Hill’s approach was executed across all overlapping grid squares re-surveyed in the second campaign (n=112) for all shared species (n=75) (available via Figshare repository, see section 7). By evaluating species occurrences relative to an 8-neighbor spatial contiguity matrix (k=8), this method standardizes recording effort by benchmarking local presence against regional occupancy probability, effectively removing duration-related sampling artifacts. Spatial dependence between survey periods was evaluated using a Mantel test with 999 permutations and linear regression models. The analysis confirmed that the spatial structure of species local frequencies was consistent between survey periods (Mantel r=0.067, p=0.0680; Figure S1B, Supplementary Material). Furthermore, the low linear determination (R^2^=0.030, p=0.0680) indicates that local variation in species occurrence reflects genuine ecological dynamics and community turnover over the 20-year span, rather than a systematic artifact of differing sampling effort duration.

Due to the lack of evidence of seasonal-based differences in the species occurrences and local distribution in 2000, records gathered for each season were merged, as in the previous period. A total of 19 visits were performed between November 2002 and July 2003.

### 2.3. Data analysis

The raw data were tabulated in four matrices of sites (squares)-by-species (112x100) and sites-by-habitat type (112x9), two for each study period, and analysed according to the needs underlying each research aim (original data and R codes are available on Figshare repository, see section 7). A new matrix with the ecological “traits” for each species was added exclusively for the 4cp analysis (see section 2.3.2). All analysis were performed with the R program (R Core Team, 2019), using the specific packages mentioned in the text.

We used a multi-scale, sequential approach in the analysis, composed of five interconnected steps, ranging from broad regional patterns to local (potential) causal mechanisms. First (step 1), a global landscape analysis (ie., the 231 squares) was conducted to detect changes in the area (ha) and proportion of change of the selected landscape classes between 1980 and 2000. This step was necessary to establish whether a net temporal trend of habitat modification indeed occurred at the regional level between the two study periods. Then (step 2) we proceed to identify the major axes of landscape change using a PCA analysis to reduce environmental dimensionality and provide synthetic, uncorrelated landscape predictors representing overall habitat trajectories (agricultural intensification vs. land abandonment through natural succession). With this general picture firmly established, we moved to the analysis of the response of the local communities to the aforementioned changes, using only the surveyed sub-sample of 112 squares. We started by presenting a global picture of the broad structural changes beyond overall species richness (step 3). To that effect, species were categorized into functional/ecological categories (phenology, biogeographic origin, habitat, diet), and the number of each category was compared between the two periods, to detect functional replacements and identify ecological “winners” and “losers” across the study area. We then proceed to evaluate species-specific shifts in local occupancy across squares (step 4) using Analysis of Deviance (dDEV) in occupancy models. This allowed us to quantify statistically significant expansions, contractions, or stability spatial distribution for individual species between the two survey periods in the area sampled. Finally (step 5), we applied a Fourth-Corner approach (Brown et al., 2014) to investigate the mechanistic drivers underlying species distribution patterns by directly linking species trait matrices, local occurrence matrices, and environmental PCA axes. This analysis identified which specific ecological traits (phenology, biogeographic origin, habitat, and diet guilds) drove species vulnerability or resilience to land-use change through environmental filtering.

#### 2.3.1. Changes in land-use between 1980 and 2000

We were not able to recover the original aerial photographs used by Ferrand de Almeida (unpublished). Therefore, to quantify land-use changes between atlas periods, we recovered the original thematic paper map (Figure S2D, Supplementary Material) produced via photointerpretation and field validation (Ferrand de Almeida unpublished). This map was then georeferenced and manually digitized in a GIS framework (QGIS 2.18). Land cover for the 2000 period was derived using the same photo-interpretation protocol on high-resolution aerial imagery available for that period (Instituto Geográfico Português, 1994). No automated classification algorithms were applied; instead, both datasets relied on direct expert interpretation and field validation. To prevent methodological artifacts, typologies from both periods were harmonized into a standardized land-cover classification scheme at equivalent spatial resolution (Figure 1B; see also Table 1).

The absolute area of each land-use category (in ha) was determined across all 231 grid squares for 1980 and 2000, with net shifts per square calculated as Δ_Area_=Area_2000_−Area_1980_. Landscape dynamics between the two study periods (ie., changes in proportional coverage) were evaluated at two hierarchical levels: (i) broad macro-classes (“by major landscape unit” in Table 1), aggregating categories into “agricultural” landscapes and “forest” (tree and shrub cover) landscapes; and (ii) individual land-use categories (“by land-use unit” in Table 1), with rare or ecologically similar types grouped prior to analysis (e.g., low and tall shrublands). Differences in proportional coverage between periods for both individual categories and aggregated macro-classes, based on the raw data of Table 1, were tested using two-sample Binomial tests for proportions (Siegel & Castellan, 1988),

Additionally, we extracted the Utilized Agricultural Area (UAA) for the Douro Wine Demarcated Region in 1980 and 1999 from the official statistical reports (INE, 1989, 1999; Figure 1B). This metric is defined as the farm area (in ha) comprising arable land (both in the open and under forest/woodland canopy), kitchen gardens, permanent crops, and permanent pastures (INE, 2009), and was used here strictly to frame the broader regional context of land-use change.

Following Chamberlain and Fuller (2000), we identified the primary trajectories of land-use change between 1980 and 2000 by applying a Principal Component Analysis (PCA) to the matrix of raw area differences (ΔArea, in hectares) between periods across all grid squares. This approach solves the potential collinearity issues inherent to co-dependent land-cover changes within bounded spatial units (squares) while providing an integrative interpretation of landscape shifts. Principal axes were derived from the correlation matrix after centering and standardizing the data. Only axes with eigenvalues >1 were retained to characterize landscape dynamics (Table 1), with loadings ≥∣0.50∣ considered dominant drivers for axis interpretation. Although all four axes with eigenvalues >1 were retained to fully describe regional landscape shifts, subsequent functional trait modeling (Fourth-Corner analysis, 4cp) focused exclusively on the first two axes (PC1 and PC2). These primary axes captured the overarching macro-regional gradients (46.7% of total landscape variance) and displayed strong spatial structure across the sampling grid (Figure S1A, Supplementary Material). In contrast, secondary axes (PC3 and PC4) represented localized, residual habitat trade-offs with negligible trait–environment correlations and were omitted from community-level inferences to prevent over-interpreting localized landscape noise. The analysis was performed using the R package multiv 1.1-6 (Murtagh, 2003).

#### 2.3.2. Changes in the bird communities between 1980 and 2000

##### Species richness and ecological structure of the assemblage

Overall, changes in species richness and selected biological and ecological categories between 1980 and 2000 were evaluated at the regional scale by counting the absolute number of species per category pooled across all grid cells (n=112 squares), according to the: (1) <u>phenological state</u> in the study area (Ferrand de Almeida, unpublished); (2) <u>biogeographic origin</u> (Suárez-Seoane et al., 2002); (3) <u>suitable/preferred habitat type (Ferrand de Almeida, unpublished)</u> and, (4) <u>diet</u> (Catry et al., 2010). Table 2 summarizes these category levels (for full details see Table S2, Supplementary Material). Absolute species counts provided a direct, zero-tolerant measure of richness shifts, avoiding artifacts associated with ratio transformations. Significant differences in species distributions across categories between periods were assessed using chi-square (χ^2^) contingency table tests at p<0.05

**Table 2.**
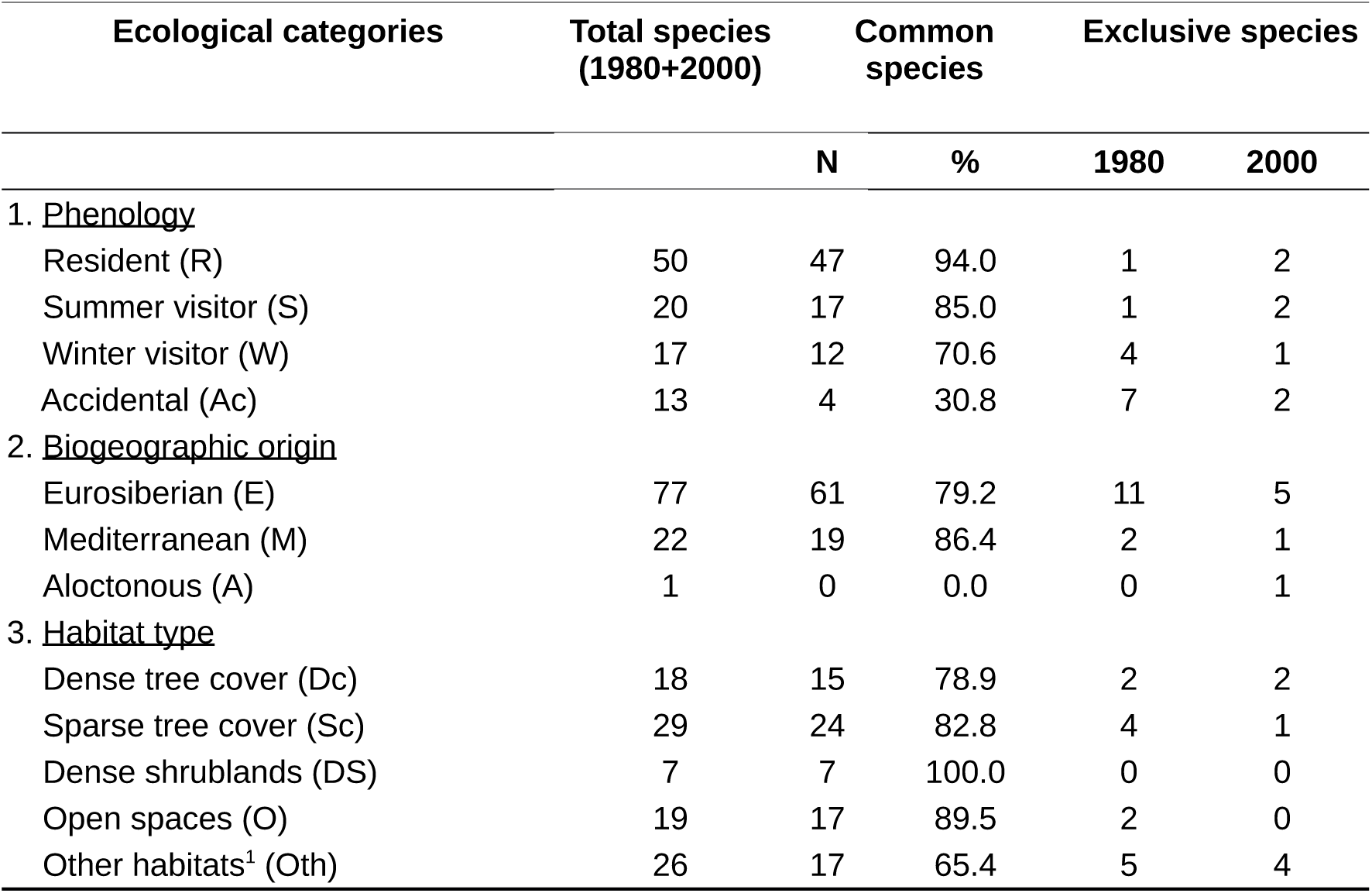
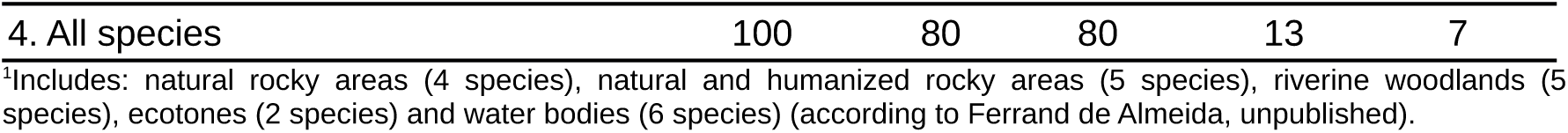
General characterization of the bird assemblages in the study area in 1980 and 2000 (n=112 squares) in regard to phenology, biogeographic origin and habitat type (see text for details). The total number of species (1980+2000), the number and percentage (%) of shared species and the number of species exclusive of each period and category are shown (see Table S1 and table S2, supplementary material, for details on the species identities and on each category, respectively). Abbreviations as in the text.

##### Changes in composition (community heterogeneity)

Changes in community composition between 1980 and 2000 were assessed using the model-based deviance analysis (statistic D) based on Generalized Linear Model (GLM) fits. This metric evaluates the species-level heterogeneity of occurrence (D_i_) across sampling units (Baeten et al., 2014). A low value of (D_i_) indicates a surplus of species that are either ‘rare’ (present in less than half of the sites) or ‘prevalent’ (present in half or more of the sites), thereby contributing little to variation within or between communities. The temporal variation in this metric, within and across sites (ΔD_i_) provides a quantitative measure of the degree of community homogenization over time (Baeten et al., 2014). Specifically, biotic homogenization increases (ΔD_i_<0) if previously common (prevalent) species become rare or if certain species become predominant (more prevalent) across communities; species contributing to ΔD_i_<0 are defined as ‘community-convergence species’. Conversely, biotic diversification increases (ΔD_i_>0) if more species become more evenly distributed among sites; species contributing to Δ ΔD_i_>0 are defined as ‘community-divergence species’ (Baeten et al., 2014). The main advantage of statistic D over traditional aggregate indexes of species heterogeneity (e.g., Shannon’s H or Jaccard dissimilarity) is that it pinpoints individual species’ contributions to the overall community shifts, allowing to disentangle homogenizing effects driven by spreading prevalent species from those caused by species losses (Baeten et al., 2014).

We calculated the species ΔD_i_’s and the community ΔD for both periods, using each sampled square as an individual “site” (A. Baeten, pers. comm.). The ‘dDEV’ R script, provided as supplementary material in Baeten et al. (2014) was used as a basis to calculate both metrics and their associated (permutated) p-values. As suggested by Baeten et al. (2014), we used the False Discovery Rate (FDR) method (Pike, 2011) to correct for multiple comparisons. The calculation of the FDR-adjusted p-values was performed with the function ‘p.adjust’, of the R base library ‘stats’ (R Core Team, 2019). The modified dDEV R script (which also includes the associated FDR values) can be found in the Figshare repository (see section 7).

##### Response of the local farmland birds to the gradients of landscape changes

To evaluate how species biological and ecological traits mediate their responses to landscape change gradients, we applied a Fourth-Corner (4cp) model-based approach using the traitglm function in the R package mvabund (Wang et al., 2012; Brown et al., 2014). The Fourth-Corner approach directly links a matrix of species occurrences across sites (L) to environmental predictors (Q, here the primary PCA landscape change axes) and species traits (R, here the biological and ecological metrics presented in Table 2). Conceptually, the core of the Fourth-Corner framework is the trait-by-environment interaction term (R×Q), which tests whether species with specific traits systematically respond positively or negatively to particular landscape change trajectories. To evaluate the predictive power of traits against individual species responses, we fitted two competing Generalized Linear Models (GLMs): (a) the “full” Fourth-Corner Model, which evaluates how ecological traits mediate responses to landscape shifts via the trait-by-environment interaction term (R×Q), while incorporating main environmental effects, species baseline terms (i.e., SpID) and a residual species-environment term (Q×SpID) to account for species-to-species variation not captured by traits (Model 3 in Brown et al., 2014); and (b) the “reduced” ‘Species-Only’ Model, which excludes trait interactions and fits separate environmental conditions for each species via the species-by-environment interaction term (Q×SpID). This model treats species themselves as traits, serving as a baseline equivalent to a classical Species Distribution Model (SDM) framework (Model 2 in Brown et al., 2014). Both models incorporated a LASSO penalty (Tibshirani, 1996) for automated variable selection and regularization. The LASSO shrinks non-informative interaction terms to zero, yielding a parsimonious model that highlights only the most robust associations (Brown et al., 2014). Models were evaluated using cross-validation (cv.glm1path), running 20 iterations with a 50:50 training/test split to select the model path that minimized predictive log-likelihood. In this regularized GLM framework, model superiority (in this case between Model 2 and Model 3) is evaluated based on the log-likelihood via deviance reduction (ΔDeviance) and BIC minimization rather than conventional R^2^ metrics.

The analysis was applied to the 2000 bird community data across n=112 grid cells, restricted to 53 focal landbird species (predominantly passerines and alike) based on three explicit criteria: (i) ecological suitability (by including species whose home ranges and resource-use patterns match the spatial resolution of the sampling units); (ii) habitat affinity (by focusing on terrestrial species directly dependent on vegetation structure and land-use practices rather than obligate wetland/coastal taxa); and (iii) statistical robustness (by retaining only species with >4 occurrences across the grid to prevent model instability driven by extreme rarity).

## 3. Results

### 3.1. Changes in land-use between 1980 and 2000

The study area remained predominantly an agricultural landscape across the evaluated time span. At the broader administrative scale of the Douro Wine Demarcated Region, the Utilized Agricultural Area exhibited considerable fluctuations between 1980 and 1999 (INE, 1989, 1999; Figure 1A, inset). However, in the two municipalities bounding the study area (Alijó and Carrazeda de Ansiães), agricultural land use remained relatively stable, with little losses or even small gains (Figure 1A).

Within our specific study grid, agricultural land cover contracted by only 7.4% (207 ha lost), whereas non-agricultural cover (encompassing shrublands, woodlands, and burnt areas) increased by approximately 5.0% (112 ha gained) (Table 1). Net changes (Δ%) of individual land-use types, showed that changes were statistically significant (p < 0.001) at both scales of evaluation (by “landcape unit” or by “ land-use type”, see section 2.3.1), though their magnitude varied markedly. Shifts were moderate when expressed relative to the total area of the respective landscape unit (range: -26.0 to +27.9), but substantially stronger when evaluating the relative turnover within specific land-use types (range: -81.6 to +100.0). The close correspondence between land-cover gains and losses across several categories strongly suggested distinct, structured trajectories of spatial transformation across the landscape.

The Principal Component Analysis (PCA) yielded four axes with eigenvalues >1, accounting for a cumulative variance of 77.1% (Table 1). PC1 axis (24.8% of variance) is a measure of agricultural restructuring, reflecting a major trajectory within active farming areas, by opposing traditional vineyards (OV, loading =+0.779, which suffered a reduction of 728 ha) and traditional olive orchards (Oli, loading =−0.528) to the expansion of intensive, fully or partially mechanized vineyards (Vin, loading =−0.824, which increased by 781 ha). Shrubland encroachment in abandoned agricultural plots also loaded positively on this axis (Shr, loading =+0.628). PC2 (21.9% of variance) is a reflection of non-agricultural land abandonment and succession. It captured both disturbance and natural succession dynamics in forestry and semi-natural areas, opposing pinewood production forests (Pin, loading =−0.664) against burnt areas (Bur, loading =+0.820) and shrubland expansion (Shr, loading =−0.527). PC3 (18.5% of variance) represents localized forest and mixed land cover trade-offs, like minor trade-offs between cork-oak woodlands (Cor, loading =+0.626) and small garden cultures (Cul, loading =−0.502), partly reflecting local transitions following wildfire events and minor agricultural shifts. Finally, PC4 (11.9% of variance) can be interpreted as a pine encroachment in open areas, depicting specific local transitions between production pinewoods (Pin, loading =+0.658) and a wet meadow (Mea, loading =−0.540), likely associated with pine regrowth in a particular open area following the cessation of traditional livestock grazing.

To better visualize the spatial patterns of landscape transformation described above, the individual PC scores were mapped across the study area (Figure S3, Supplementary Material). These maps clearly point out the contrasting paths of spatial dynamics of landscape change captured by each PCA axis.

### 3.2 Changes in bird assemblages between 1980 and 2000

#### Species composition and ecological structure (ecological guilds)

Overall, 100 species were recorded for the two periods considered, of which 80 occurred regularly (Table 2; see also Table S2, Supplementary Material, for details). The majority of the recorded species (77%) had an Eurosiberian origin, the rest being Mediterranean (22%) or allochthonous (African) origin (1%). The resident species formed half of the community, and 37% were found to be migratory species (both spring and winter visitors). Approximately half of the species (48%) were characteristic of open habitats, either from natural (29%) or farmland (19%) type (categories “sparse tree cover” (“Sc”) and “open” (“O”), respectively), 25% were forest generalists (*sensu latu*: woodlands (18%) and shrublands (7%), categories “dense tree cover” (“Dc”) and “dense shrublands” (“DS”), Table 2), and 26% of the species used other habitats.

Shifts in regional community composition between 1980 and 2000 was negligible, with 80% of the species common to both periods (Table 2). Also, there was no significant evidence of temporal changes in the global composition of the community according to biogeographic origin (χ^2^=1.29, df=2, p<0.53), phenology (χ^2^=7.62, df=3, p>0.06) or habitat (χ^2^=0.15, df=4, p<0.99).

#### Community heterogeneity and species-specific patterns

The bird community in the study area did not show significant evidence of biotic homogenization between 1980 and 2000 (ΔD=-99.359, p<0.48, n_sites_=112, n_species_=100), but species-specific significant patterns were found. In all, 30 species presented meaningful changes in their deviances (ΔD_i_’s). Of these, 17 (57%) were ‘community-convergent’ species, while 13 (43%) were “community-divergent” species (Table 3).

**Table 3.**
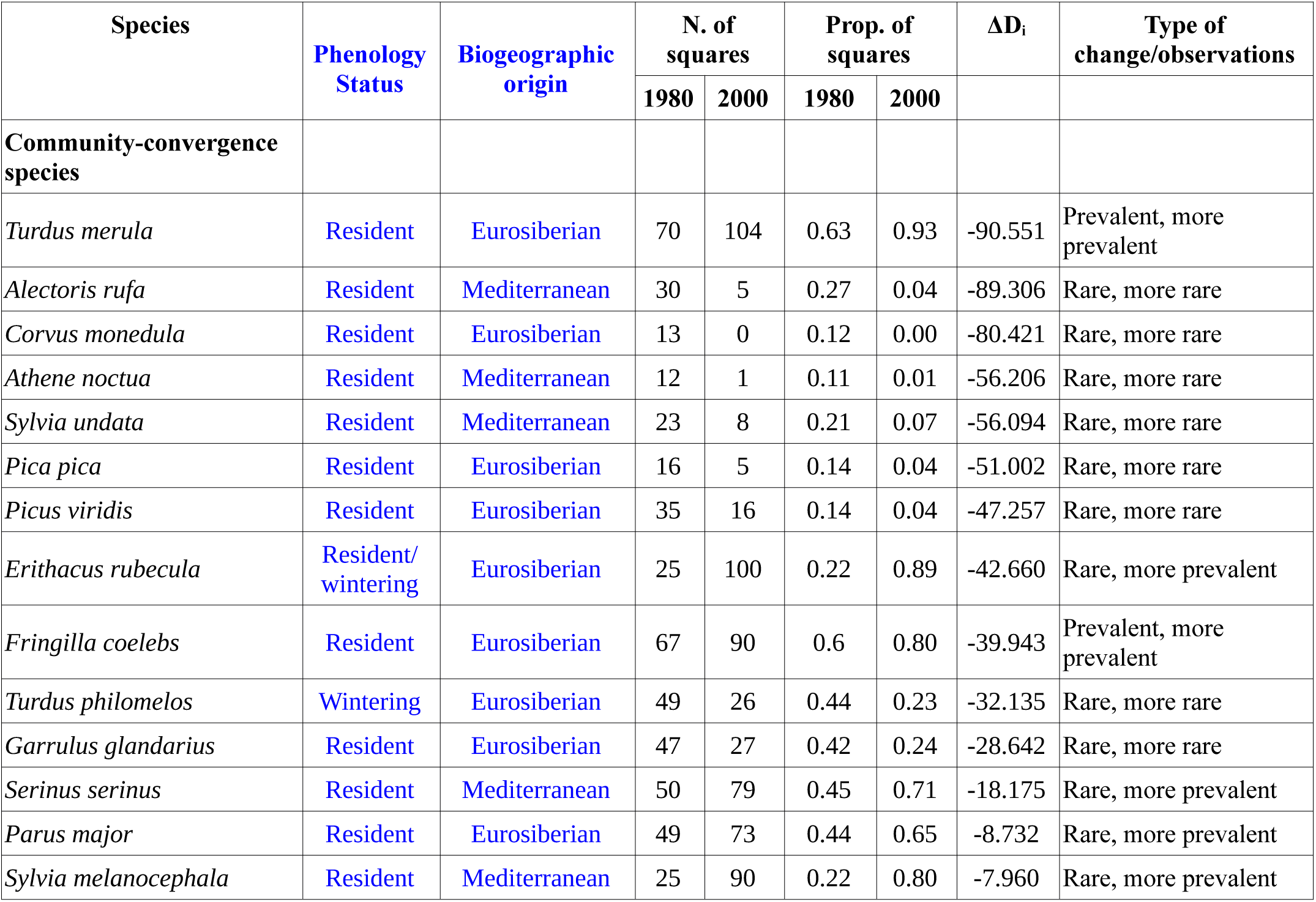
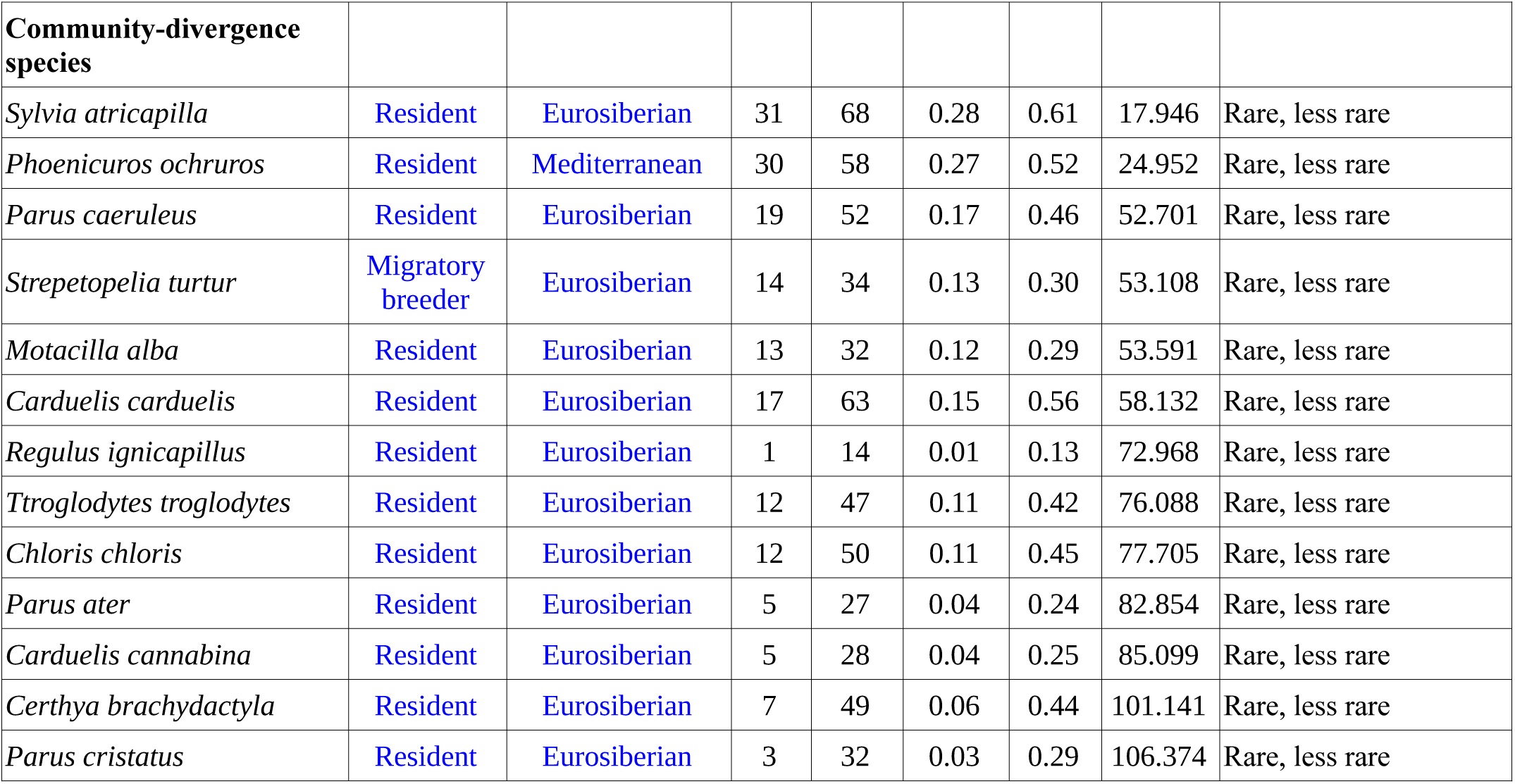
Species contributing significantly to the compositional heterogeneity of the community between 1980 and 2000. (**ΔD_i_**, p<0.05, adjusted for False Discovery Rate, FDR) The direction of change in occurrence is categorized based on initial species rarity in 1980 (where “rare” species occurred in <56 grid cells [half of the total] and “prevalent” species in ≥56 cells) and their temporal trajectory between 1980 and 2000 (e.g., “Rare, more prevalent” indicates a species that was rare in 1980 but expanded its occurrence by 2000). See Table S1 in Supplementary Material for the complete list of all species, **ΔD_i_** values, and associated p-values.

| Species | Phenology Status | Biogeographic origin | N. of squares | | Prop. of squares | | $\Delta D_i$ | Type of change/observations |
| --- | --- | --- | --- | --- | --- | --- | --- | --- |
|  |  |  | 1980 | 2000 | 1980 | 2000 |  |  |
| **Community-convergence species** |  |  |  |  |  |  |  |  |
| *Turdus merula* | Resident | Eurosiberian | 70 | 104 | 0.63 | 0.93 | -90.551 | Prevalent, more prevalent |
| *Alectoris rufa* | Resident | Mediterranean | 30 | 5 | 0.27 | 0.04 | -89.306 | Rare, more rare |
| *Corvus monedula* | Resident | Eurosiberian | 13 | 0 | 0.12 | 0.00 | -80.421 | Rare, more rare |
| *Athene noctua* | Resident | Mediterranean | 12 | 1 | 0.11 | 0.01 | -56.206 | Rare, more rare |
| *Sylvia undata* | Resident | Mediterranean | 23 | 8 | 0.21 | 0.07 | -56.094 | Rare, more rare |
| *Pica pica* | Resident | Eurosiberian | 16 | 5 | 0.14 | 0.04 | -51.002 | Rare, more rare |
| *Picus viridis* | Resident | Eurosiberian | 35 | 16 | 0.14 | 0.04 | -47.257 | Rare, more rare |
| *Erithacus rubecula* | Resident/<br>wintering | Eurosiberian | 25 | 100 | 0.22 | 0.89 | -42.660 | Rare, more prevalent |
| *Fringilla coelebs* | Resident | Eurosiberian | 67 | 90 | 0.6 | 0.80 | -39.943 | Prevalent, more prevalent |
| *Turdus philomelos* | Wintering | Eurosiberian | 49 | 26 | 0.44 | 0.23 | -32.135 | Rare, more rare |
| *Garrulus glandarius* | Resident | Eurosiberian | 47 | 27 | 0.42 | 0.24 | -28.642 | Rare, more rare |
| *Serinus serinus* | Resident | Mediterranean | 50 | 79 | 0.45 | 0.71 | -18.175 | Rare, more prevalent |
| *Parus major* | Resident | Eurosiberian | 49 | 73 | 0.44 | 0.65 | -8.732 | Rare, more prevalent |
| *Sylvia melanocephala* | Resident | Mediterranean | 25 | 90 | 0.22 | 0.80 | -7.960 | Rare, more prevalent |

| Community-divergence species |  |  |  |  |  |  |  |  |
| --- | --- | --- | --- | --- | --- | --- | --- | --- |
| *Sylvia atricapilla* | Resident | Eurosiberian | 31 | 68 | 0.28 | 0.61 | 17.946 | Rare, less rare |
| *Phoenicurus ochruros* | Resident | Mediterranean | 30 | 58 | 0.27 | 0.52 | 24.952 | Rare, less rare |
| *Parus caeruleus* | Resident | Eurosiberian | 19 | 52 | 0.17 | 0.46 | 52.701 | Rare, less rare |
| *Streptopelia turtur* | Migratory breeder | Eurosiberian | 14 | 34 | 0.13 | 0.30 | 53.108 | Rare, less rare |
| *Motacilla alba* | Resident | Eurosiberian | 13 | 32 | 0.12 | 0.29 | 53.591 | Rare, less rare |
| *Carduelis carduelis* | Resident | Eurosiberian | 17 | 63 | 0.15 | 0.56 | 58.132 | Rare, less rare |
| *Regulus ignicapillus* | Resident | Eurosiberian | 1 | 14 | 0.01 | 0.13 | 72.968 | Rare, less rare |
| *Troglodytes troglodytes* | Resident | Eurosiberian | 12 | 47 | 0.11 | 0.42 | 76.088 | Rare, less rare |
| *Chloris chloris* | Resident | Eurosiberian | 12 | 50 | 0.11 | 0.45 | 77.705 | Rare, less rare |
| *Parus ater* | Resident | Eurosiberian | 5 | 27 | 0.04 | 0.24 | 82.854 | Rare, less rare |
| *Carduelis cannabina* | Resident | Eurosiberian | 5 | 28 | 0.04 | 0.25 | 85.099 | Rare, less rare |
| *Certhya brachydactyla* | Resident | Eurosiberian | 7 | 49 | 0.06 | 0.44 | 101.141 | Rare, less rare |
| *Parus cristatus* | Resident | Eurosiberian | 3 | 32 | 0.03 | 0.29 | 106.374 | Rare, less rare |

The ‘community-convergent’ group was found to be rather heterogeneous. It included rare species that became even rarer, from several taxonomic origins and (mostly) from open/mixed habitats. (eg., *Alectoris rufa*, *Corvus monedula*, which actually gone extinct from the area), *Athene noctua, Sylvia undata* or *Picus sharpei)*, but also prevalent species that became more prevalent (eg., *Turdus merula* and *Fringilla coelebs),* and also species that were rare in 1980 but became prevalent in 2000 (*Serinus serinus*, *Parus major*, *Sylvia melanocephala* and, particularly, *Erithacus rubecula*, the only confirmed case of a species that altered its phenology status in the study period, from a strict winter visitor (Ferrand de Almeida, unpublished) to resident.

The ‘community-divergent’ species, on the other hand, were all ‘rare’ passerine species that became less rare, and belonged to two main habitat/feeding guilds: strict woodland/shrubland species (eg., *Sylvia atricapilla*, *Troglodytes troglodytes*, *Lophophanes cristatus*, *Periparus ater*, *Certhia brachydactyla*, *Regulus ignicapillus)*, and open habitat species *sensu latu* (ie. categories “open spaces” (“O”) and “sparse tree cover” (“Sc”) of Table 2) (e.g., *Phoenicurs ochrurus, Motacilla alba, Carduelis spp., Streptopelia turtur* and *Chloris chloris)*.

#### Species-specific and ecological traits of the local birds associated to the major drivers of landscape change

The model with the best predictive performance was the ‘species model’ (Model 2): LogLik=−2591.17 and Deviance=5182.34 *versus* LogLik=−2622.74 and Deviance=5245.53 of the ‘traits model’ (Model 3).

The results of the ‘traits model’ showed that habitat type, diet, and biogeographic origin, in this order, were the ecological traits that best explained bird associations with the axes of landscape change, whereas phenology was not relevant (Figure 2, left). The strongest associations were found along PC1 axis (the direction of change towards agricultural restruturing and localized shrubland encroachment). The traits associated with open or semi-open habitats (“HabitatO” and “HabitatSc” categories) correlated positively with this axis. Conversely, habitat traits associated with dense canopy cover (“`HabitatDc”`), showed a strong negative correlation with PC1. The agricultural restruturing was also detrimental (ie,. correlated negatively) for species of omnivorous diets (“DietOmn”) and with a large distribution (Eurosiberian origin, “OriginE”). Along PC2 (non-agricultural land abandonment and succession), all associations were restricted to the habitat guild. Strong negative correlations were found for open habitats (“`HabitatO”`) and other habitat types (rocky areas, aquatic habitats) (“HabitatOth”), while a weaker positive correlation was observed with dense cover habitats (“HabitatDc”). Functional trait associations along PC3 and PC4 were considerably weaker and more diffuse, and thus were not examined here (Figure 2, left).

**Figure 2.**
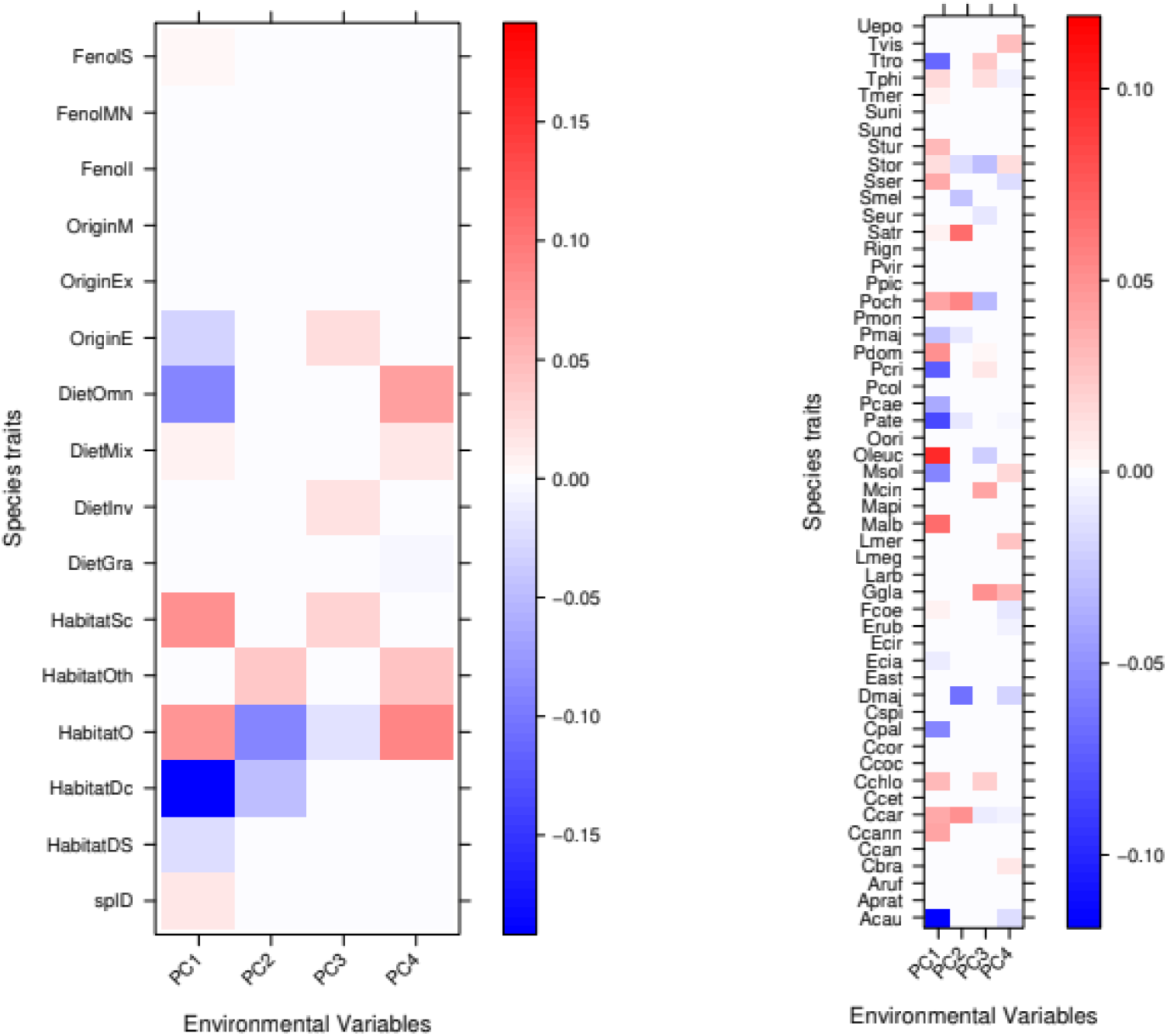
Graphic results of a fourth-corner analysis for the two models examined: (a) the variant of the traditional (environment*traits) model, which includes a species-specific identification term (spID) (model 3 of Brown et al., 2014), and (b) the “species-only” (i.e., environment * species) model where the trait matrix is forfeited and the species themselves are used as response (model 2) The colour gradient represents the magnitude (strength) of the coefficients retained in the model for each term (ie., pair environmental variable - response variable), from negative (blue) to positive (red) association; the darker the tone, the higher the value of the coefficient. Non-significant terms are left white. Species traits variables (model 3) are: **Fenol** - Phenology (’S’: Sedentary, T: Wintering; ‘MN’: Breeding); **Origin** (‘M’: Mediterranean, ‘Ex’: Exotic (alloctonous), ‘E’: Eurosiberian); **Diet** (’Omn’: Omnivorous; ‘Mix’: Mixed; ‘Inv’: Invertebrates; ‘Gran’: Granivorous); **Habitat** (‘Sc’: Sparse cover; ‘Oth’: Other; ‘O’: Open; Dc’: Dense cover; ‘DS’: Dense shrubland); spID - Species-specific indentification. Species names codes (model 2) as in Table Si. See text for details.

In the ‘species model’, a total of 32 species (out of 53) responded significantly to one or more axes of landscape change (Figure 2, right). The highest number of species (23) and the strongest correlations were associated with PC1, either positively (e.g., *Oenanthe leucura*) or negatively (e.g., *Troglodytes troglodytes*, *Lophophanes cristatus*, *Periparus ater*, *Aegithalos caudatus*, *Columba palumbus*, and *Monticola solitarius*). In contrast, PC2 captured responses driven by non-agricultural land abandonment and fire disturbance, with species sorting according to habitat openness; open-habitat and scrub specialists (such as *Phoenicurus ochruros* and *Sylvia atricapilla*, respectively) were positively associated with this axis, whereas forest-dependent species (e.g., *Dendrocopos major*) exhibited negative associations. Responses along the remaining axes involved a bit more species (PC3: 11 species; PC4: 13 species), but had generally weaker correlations (Figure 2, right), thus being of less interpretative value.

## 4. Discussion

Land-use change is one of the major drivers of change impacting biodiversity in the Anthropocene (Newbold et al., 2015; Regos et al., 2016). Habitat destruction or landscape simplification is a major consequence of both intensification and land abandonment dynamics (Gámez-Virués et al., 2015). At the community level, these processes typically lead to biotic homogenization, where specialist species are replaced by widespread generalists (Clavel et al., 2011; Reino et al., 2018). However, most studies focus solely on either intensification or abandonment, failing to provide a holistic overview of their simultaneous or synergistic impacts on bird assemblages (see introduction). Here, we combined historical (1980) and more recent (2000) data on both bird communities and land-use patterns to evaluate long-term response patterns in a Mediterranean landscape where both drivers co-occurred over two decades. In this context, our findings demonstrate that the primary trajectories of land-use change identified in the region (agricultural restructuring and land abandonment) were reflected at both species-specific and functional-trait levels but did not translate into broad-scale structural shifts at the community level. This suggests that local avian assemblages possessed sufficient ecological resilience to buffer the potencially synergistic dynamics of these two contrasting drivers.

In fact, the analysis of land-use trends between 1980 and 2000 suggests that the study area remained essentially a traditional farmland mosaic, with less than 10% of agricultural land converted to other uses. Nevertheless, clear trajectories of structural change were identified (Table 1; Figure S3 in Supplementary Material). While PC1 (agricultural restructuring) captured the transition from traditional vineyards to intensified or semi-intensified vineyards alongside localized shrubland encroachment, PC2 (non-agricultural land abandonment and succession) depicted forestry dynamics and disturbance, opposing production pine stands against burnt areas and early-successional shrublands. Overall, the competing spatial pressures of these two main drivers counteracted each other at the landscape scale, preventing the emergence of a completely homogenized landscape and maintaining a heterogeneous mosaic until the early 2000s. This structural persistence contrasts with the more widespread agricultural abandonment reported in other northern Portuguese farmlands (Moreira et al., 2001; Azevedo et al., 2011). Maintaining landscape-level heterogeneity allowed the avian community to preserve its initial, diversified species pool (a mixture of resident farmland and woodland generalists with broad biogeographical distributions). This structural diversity is fundamental to sustaining ecosystem functional diversity and resilience against environmental shifts (Mori et al., 2013), explaining why local bird assemblages remained remarkably stable in taxonomic composition. Indeed, between 1980 and 2000 only two species (*Neophron percnopterus* and *Corvus monedula*) became locally extinct and, in both cases, this mirrors broader national trends (Equipa Atlas, 2008) and/or (in the case of *N. percnopterus*) the loss of suitable nesting habitats due to potential disturbance (Ferrand de Almeida, *own obs.*), rather than localized landscape degradation. This community-level stability is further supported by the absence of significant biotic homogenization (ΔD<0). At the intra-guild and species-specific levels, however, community responses were more distinct and aligned with functional predictions.

The effects of agricultural restructuring (PC1), typically associated with vegetation simplification and a corresponding loss of species diversity (Stoate et al., 2009), were strongly structured by habitat cover, diet, and biogeographical origin (Forth-Corner model 3). In contrast, non-agricultural land abandonment (PC2) exhibited a narrower functional footprint, being restricted primarily to habitat-use guilds. Regarding biogeographical origin, Eurosiberian species exhibited strong negative associations with PC1. These are mainly broadly distributed woodland species associated with closed canopy cover (e.g., *Troglodytes troglodytes*, *Lophophanes cristatus*, *Periparus ater, Aegithalos caudatus, Columba palumbus, and Monticola solitarius*), which avoided restructured, open agricultural areas. This aligns with Suárez-Seoane et al. (2002), who reported a strong preference among Eurosiberian birds for dense, closed vegetation structures, particularly during the breeding season. The dominance of PC1 as a driver of community reassembly was confirmed at the species level (model 2 of the Fourth-Corner analysis), where 23 species responded significantly along this axis, compared to 8 species along PC2. Individual species responses further illustrated the nuanced ecological pressures operating across the landscape. For instance, *Periparus ater*, a woodland specialist, was negatively associated with PC1, reflecting the loss or degradation of suitable tree cover within agricultural areas. Conversely, *Oenanthe leucura*, an open-habitat specialist dependent on traditional vineyard terraces and olive orchards (Múrias et al., 2008), correlated positively with PC1, likely responding to increased open ground availability resulting from agricultural modernization and localized clearing. The higher explanatory power of species-specific models compared to trait-only models suggests that species identity plays a stronger role in driving community responses to landscape change than the functional traits evaluated here, likely reflecting species-level ecological nuances or unmeasured traits.

In short, the results of our study suggest that the guild structure and composition of the bird communities were not strongly associated with the co-occurrence of intensification and land abandonment dynamics. Causes underlying our results may include the combination of (a) an assemblage of resilient species prior to the onset of the land-use change dynamics with (b) the buffering effect of the two competing drivers of land use change.

A critical challenge when comparing multi-decadal atlas datasets stems from variations in survey duration and recording effort (in the present case, six years in 1979–1984 vs. two years in 2002–2003). However, the concentrated, high-intensity protocol implemented during the second atlas was explicitly designed to rapidly capture local species composition within each grid square. In grid-based occurrence surveys, spatial effort standardization across neighboring units is the primary determinant of comparability, rather than temporal duration per se (Hill, 2012). Formal validation using Hill’s neighborhood local frequency index confirmed that spatial occurrence structures were consistent between periods across the continuously monitored grid squares (n=112; Mantel r=0.067,p=0.0680; R^2^=0.030), preventing duration-driven reporting artifacts. While distribution-only data cannot capture subtle abundance fluctuations (Plieninger et al., 2016; Ehrlén and Morris, 2015), an inherent limitation of “first-generation” atlases (Keller, 2017), this neighborhood-based effort validation ensures that observed changes reflect genuine community-level dynamics rather than sampling artifacts.

Our findings also provide key historical insights by (1) evaluating the potentially synergistic impacts of agricultural restructuring and land abandonment at regional scales (Sanderson et al., 2013; Katayama et al., 2015); (b) by extending long-term ecological assessments in northern Portugal (Santana et al., 2014; Silva et al., 2018) back to the pre-1990 baseline; and (c) by establishing a socio-ecological reference situation prior to the major Common Agricultural Policy (CAP) reforms of the 1990s in Portugal (Jones et al., 2011). Indeed, an added value of this study lies in the recovery of an old atlas dataset illustrating the baseline state of an avian community within a farmland mosaic of NE Portugal prior to the major landscape transformations following the 1993 CAP reform.

In summary, this study provides empirical evidence of the potential impacts arising from the spatio-temporal co-occurrence of agricultural intensification and abandonment in recent decades (e.g., Sanderson et al., 2013; Katayama et al., 2015; Aune et al., 2018). Although the interaction between these dual drivers has received limited attention until recently, its relevance is expected to increase across European landscapes (e.g., Katayama et al., 2015). Indeed, contemporary land-use trends in Europe increasingly foster land abandonment and subsequent natural succession, creating spatial configurations where agricultural intensification and rewilding processes interact within shared geographic contexts. More broadly, this work underscores that even spatially restricted, presence/absence atlas datasets can still offer substantial ecological value when addressing complex global change issues. By employing appropriate modern analytical frameworks, such historical records provide vital baselines, particularly if the original data is duly curated and made publicly available, as in the present case (the data are available on the Figshare repository). We hope these findings encourage the publication, curation, and open sharing of similar legacy datasets.

Finally, beyond its theoretical value, this dataset provides a crucial pre-construction benchmark prior to the major infrastructure development of the Foz-Tua Dam (initiated in 2006 and completed in 2017; Profico, 2008). As the resulting reservoir flooded extensive valley habitats across the study region, including several in the surveyed grid squares, our spatial baseline offers an essential historical reference to evaluate how long-term landscape shifts interact with acute habitat loss in avian communities. This is particularly relevant for ongoing monitoring within the Regional Natural Park of the Tua Valley (ICNF, 2026) and as a broader case study for the highly modified Douro Demarcated Region.

## Supporting information

Includes Supplementary Figures S1-S3 and Supplementary Tables ST1 and ST2

## Acknowledgements

The authors are indebted to all people who helped in the fieldwork in 1979-1984, and provided hosting (in both periods) for this study.

## 7. Data, scripts and code availability

The original datasets and R scripts codes used in the preparation of this article are available in the Figshare repository, through the URL **xxxxx** [under embargo until the end of the revision process] An acompanying txt file with the detailed explanation of each file/script is also included.

## References

Andresen, T., De Aguiar, F.B. and Curado, M.J. (2004). The Alto Douro wine region greenway. Landscape and urban planning, 68(2-3): 289–303. doi – 10.1016/S0169-2046(03)00156-7

Aune S., Anders B., Hovstad, K.A. (2018) Loss of semi-natural grassland in a boreal landscape: impacts of agricultural intensification and abandonment, Journal of Land Use Science, 13(4), 375–390, DOI: 10.1080/1747423X.2018.1539779

Azevedo J.C., Moreira C., Castro J.P., Loureiro C. (2011). Agriculture Abandonment, Land-use Change and Fire Hazard in Mountain Landscapes in Northeastern Portugal. In: Li C., Lafortezza R., Baeten, L., Warton, D.I., Van Calster, H., De Frenne, P., Verstraeten, G., Bonte, D., Bernhardt-Römermann, M., Cornelis, J., Decocq, G., Eriksson, O. and Hedl, R. (2014). A model-based approach to studying changes in compositional heterogeneity. Methods in Ecology and Evolution, 5(2): 156–164. doi – 10.1111/2041-210X.12137

Barreto, A. (1993). *Douro*. Edições Inapa. 180 pp.

BirdLife International (2004) Birds in the European Union: a status assessment. Wageningen, The Netherlands: BirdLife International. 50 pp.

Blondel, J. (1969). Méthodes de dénombrement des populations d’oiseaux. In: Lamotte et Bourlière: Problèmes d’Ecologie: l’échantillonage des peuplements animaux des milieux terrestres, 97–151. Paris, Masson.

Bonthoux, S., Barnagaud, J.Y., Goulard, M. and Balent, G. (2013). Contrasting spatial and temporal responses of bird communities to landscape changes. Oecologia, 172(2): 563–574. doi – 0.1007/s00442-012-2498-2

Brambilla, M., Casale, F., Bergero, V., Bogliani, G., Crovetto, G.M., Falco, R., Roati, M. and Negri, I. (2010). Glorious past, uncertain present, bad future? Assessing effects of land-use changes on habitat suitability for a threatened farmland bird species. Biological Conservation, 143(11): 2770–2778. doi – 10.1016/j.biocon.2010.07.025

Brown, A.M., Warton, D.I., Andrew, N.R., Binns, M., Cassis, G. and Gibb, H. (2014). The fourth-corner solution–using predictive models to understand how species traits interact with the environment. Methods in Ecology and Evolution 5(4): 344–352. doi – 10.1111/2041-210X.12163

Catry, P., Costa, H., Elias, G. and Matias, R. (2010). Aves de Portugal: Ornitologia do Território Continental (Birds of Portugal: Ornithology of the Continental Territory). Assirio & Alvin.

Chamberlain, D.E. and Fuller, R.J. (2000). Local extinctions and changes in species richness of lowland farmland birds in England and Wales in relation to recent changes in agricultural land-use. *Agriculture*, Ecosystems & Environment 78(1): 1–17. doi – 10.1016/S0167-8809(99)00105-X

Clavel, J., Julliard, R. and Devictor, V. (2011). Worldwide decline of specialist species: toward a global functional homogenization? Frontiers in Ecology and the Environment 9(4): 222–228. doi – 10.1890/080216

De Palma, A., Sanchez-Ortiz, K., Martin, P.A., Chadwick, A., Gilbert, G., Bates, A.E., Börger, L., Contu, S., Hill, S.L. and Purvis, A. (2018). Challenges with inferring how land-use affects terrestrial biodiversity: Study design, time, space and synthesis. In Advances in Ecological Research 58: 163–199. Academic Press. doi – 10.1016/bs.aecr.2017.12.004

Donald, P.F., Green, R.E. and Heath, M.F. (2001). Agricultural intensification and the collapse of Europe’s farmland bird populations. Proceedings of the Royal Society of London B: Biological Sciences, 268(1462): 25–29. doi – 0.1098/rspb.2000.1325

Donald, P.F., Pisano, G., Rayment, M.D. and Pain, D.J. (2002). The Common Agricultural Policy, EU enlargement and the conservation of Europe’s farmland birds. Agriculture, Ecosystems & Environment 89(3): 167–182. doi – 10.1016/S0167-8809(01)00244-4

Donald, P.F., Sanderson, F.J., Burfield, I.J. and Van Bommel, F.P. (2006). Further evidence of continent-wide impacts of agricultural intensification on European farmland birds, 1990–2000. Agriculture, Ecosystems & Environment 116(3-4): 189–196. doi – 10.1016/j.agee.2006.02.007

Dunn, A.M. and Weston, M.A. (2008). A review of terrestrial bird atlases of the world and their application. Emu-Austral Ornithology 108(1): 42–67. doi – 10.1071/MU07034

Equipa Atlas (2008). *Atlas das Aves Nidificantes em Portugal (1999-2005)* (Atlas of the Breeding Birds of Portugal (1999-2005)). Instituto de Conservação da Natureza e Biodiversidade, Sociedade Portuguesa para o Estudo das Aves, Parque Natural da Madeira and Secretaria Regional do Ambiente e do Mar. Assírio & Alvim, Lisboa.

Ehrlén, J. and Morris, W.F. (2015). Predicting changes in the distribution and abundance of species under environmental change. Ecology Letters 18: 303–314.

Ferrand de Almeida N. (unpublished). Estrutura da Comunidade de Vertebrados Terrestres de um Ecossistema Agrícola (Structure of the Vertebrate Community of an Agricultural Ecosystem). Unpublished Report, 1st Prize, National Environment Year 1987. SEA, Lisbon. 318 pp.

Flynn, D.F., Gogol-Prokurat, M., Nogeire, T., Molinari, N., Richers, B.T., Lin, B.B., Simpson, N., Mayfield, M.M. and DeClerck, F. (2009). Loss of functional diversity under land use intensification across multiple taxa. Ecology Letters 12(1): 22–33. doi – 10.1111/j.1461-0248.2008.01255.x

Fonderflick, J., Caplat, P., Lovaty, F., Thévenot, M. and Prodon, R. (2010). Avifauna trends following changes in a Mediterranean upland pastoral system. *Agriculture*, Ecosystems & Environment 137(3-4): 337–347. doi – 10.1016/j.agee.2010.03.004

Gámez-Virués, S., Perović, D.J., Gossner, M.M., Börschig, C., Blüthgen, N., de Jong, H., Simons, N.K., Klein, A.-M., Krauss, J., Maier, G., Scherber, C., Steckel, J., Rothenwöhrer, C., Steffan-Dewenter, I., Weiner, C.N., Weisser, W., Werner, M., Tscharntke, T., and Westphal, C. (2015). Landscape simplification filters species traits and drives biotic homogenization. Nature Communications 6 (8568): 1–8. doi – 10.1038/ncomms9568

Gibbons, D.W., Donald, P.F., Bauer, H.G., Fornasari, L. and Dawson, I.K. (2007). Mapping avian distributions: the evolution of bird atlases. Bird Study 54(3): 324–334. doi – 10.1080/00063650709461492

Guerrero, I., Morales, M.B., Oñate, J.J., Aavik, T., Bengtsson, J., Berendse, F., Clement, L.W., Dennis, C., Eggers, S., Emmerson, M. and Fischer, C. (2011). Taxonomic and functional diversity of farmland bird communities across Europe: effects of biogeography and agricultural intensification. Biodiversity and Conservation 20(14): 3663–3681. doi – 10.1007/s10531-011-0156-3

Guilherme, J.L. and Pereira, H.M., (2013) Adaptation of bird communities to farmland abandonment in a mountain landscape. PloS one, 8(9), p.e73619. doi – 10.1371/journal.pone.0073619

ICNF (2026). Parque Natural Regional do Vale do Tua https://www.icnf.pt/conservacao/rnapareasprotegidas/reservasnaturais. Retrieved in 2026/07/01.

INE - Instituto Nacional de Estatística (2002). Censos (2001): Resultados Ddefinitivos do XIV Recenseamento Geral da população & IV Recenseamento Geral da Habitação (Censuses (2001) Final Results of the XIV General Population Census & IV General Housing Census (2001)) . Volume 2, 382 pp. Instituto Nacional de Estatística, Lisboa

INE (1989). Recenseamento Geral Agrícola 1989 (Concelhos) (1989 Agricultural Census (Counties)). Instituto Nacional de Estatística (INE), Relatório Técnico, Lisboa. 302 pp.

INE (1999). Recenseamento Geral da Agricultura 1999 (Trás-os-Montes) (1999 Agricultural Census (Trás-os-Montes)). Instituto Nacional de Estatística (INE), Relatório Técnico, Lisboa. 160 pp + anexos.

INE (2009). Indicadores Agro-Ambientais 1989-2007). Instituto Nacional de Estatística (INE), Relatório Técnico, Lisboa. 175 pp + anexos.

Herrando, S., Anton, M., Sardà-Palomera, F., Bota, G., Gregory, R.D. and Brotons, L. (2014). Indicators of the impact of land use changes using large-scale bird surveys: land abandonment in a Mediterranean region. Ecological Indicators 45: 235–244. doi – 10.1016/j.ecolind.2014.04.011

Jiguet, F., Devictor, V., Julliard, R. and Couvet, D. (2012). French citizens monitoring ordinary birds provide tools for conservation and ecological sciences. Acta Oecologica 44: 58–66. doi – 10.1016/j.actao.2011.05.00

PORDATA (2018). População residente segundo os Censos: total e por sexo. (Resident Population according to the Census: total and by sex). PORDATA – Estatísticas, gráficos e indicadores de Municípios, Portugal e Europa. Fundação Francisco Manuel dos Santos, Porto. Retrieved 2020-04-10, from http://www.pordata.pt3

Jeliazkov, A., Mimet, A., Chargé, R., Jiguet, F., Devictor, V. and Chiron, F. (2016). Impacts of agricultural intensification on bird communities: New insights from a multi-level and multi-facet approach of biodiversity. Agriculture, Ecosystems & Environment 216: 9–22. doi – 10.1016/j.agee.2015.09.017

Jones, N., De Graaff, J., Rodrigo, I. and Duarte, F. (2011). Historical review of land use changes in Portugal (before and after EU integration in 1986) and their implications for land degradation and conservation, with a focus on Centro and Alentejo regions. Applied Geography 31(3): 1036–1048. doi – 10.1016/j.apgeog.2011.01.024

Katayama, N., Osawa, T., Amano, T. and Kusumoto, Y. (2015). Are both agricultural intensification and farmland abandonment threats to biodiversity? A test with bird communities in paddy-dominated landscapes. Agriculture, Ecosystems & Environment 214: 21–30. doi – 10.1016/j.agee.2015.08.014

Keenleyside, C., Tucker, G. and McConville, A. (2010). Farmland Abandonment in the EU: an Assessment of Trends and Prospects. Institute for European Environmental Policy, London.

Keller, V. (2017). Atlases as a tool to document changes in distribution and abundance of birds. Vogelwelt 137: 43–52.

Merckx, T. and Pereira, H.M. (2015). Reshaping agri-environmental subsidies: From marginal farming to large-scale rewilding. Basic and Applied Ecology 16 (2): 95–113. doi – 10.1016/j.baae.2014.12.003

Moreira, F., Ferreira, P.G., Rego, F.C. and Bunting, S. (2001). Landscape changes and breeding bird assemblages in northwestern Portugal: the role of fire. Landscape Ecology 16(2): 175–187. doi – 10.1023/A:1011169614489

Moreira, F., Beja, P., Morgado, R., Reino, L., Gordinho, L., Delgado, A. and Borralho, R., (2005). Effects of field management and landscape context on grassland wintering birds in Southern Portugal. Agriculture, Ecosystems & Environment 109(1-2): 59–74. doi – 10.1016/j.agee.2005.02.011

Mori, A. S., Furukawa, T. and Sasaki, T. (2013). Response diversity determines the resilience of ecosystems to environmental change. Biological Revue 88: 349–364. doi – 10.1111/brv.12004

Múrias, T., Ribeiro, S.B., Nunes, A.I. and Gomes, C. (2008). On the occurrence and nesting of the Black Wheatear Oenanthe leucura in the area od Foz-Tua (Douro Wine Demarcated Region, NE Portugal). Airo 18: 34–39.

Múrias, T., Ribeiro, S.B., Lomba, A. and Ferrand de Almeida, N. (2026). Datasets and R Scripts used in the present paper. Figshare Repository, https://figshare.com/s/c6650e164d5ac897105b

Murthagh, F. (2003). Multltivariate Data Analysis Routines. R package version 1.1-6. https://cran.r-project.org/src/contrib/Archive/multiv/

Newbold, T., Lawrence, N.H., Hill, S.L.L., Contu, S., Lysenko, I., Senior, R.A., Börger, L., Bennett, D.J., Choimes, A., Collen, B., Day, J., De Palma, A., Díaz, S., Echeverria-Londoño, S., Edgar, M.J., Feldman, A., Garon, M., Harrison, M.L.K., Alhusseini, T., Ingram, D.J., Itescu, Y., Kattge, J., Kemp, V., Kirkpatrick, L., Kleyer, M., Pinto Correia, D.L., Martin, C.D., Meiri, S., Novosolov, M., Pan, Y., Phillips, H.R.P., Purves, D.W., Robinson, A., Simpson, J., Tuck, S.L., Weiher, E., White, H.J., Ewers, R.M., Mace, G.M., Scharlemann, J.P.W., Purvis, A. (2015). Global effects of land use on local terrestrial biodiversity. Nature 520: 45–50. doi – 10.1038/nature14324

Nikolov, S.C. (2010). Effects of land abandonment and changing habitat structure on avian assemblages in upland pastures of Bulgaria. Bird Conservation International 20(2): 200–213. doi – 10.1017/S0959270909990244

Parachinni, M.L., Petersen, J.E., Hoogeveen, Y., Bamps, C., Burfield, I. and van Swaay, C. (2008). High nature value farmland in Europe. An estimate of the distribution patterns on the basis of land cover and biodiversity data. Technical Report EUR, 23480.

Parodi, J.M., Cuthbert, F.J. and Decker, E.H. (2001). The effect of 50 years of landscape change on species richness and community composition. Global Ecology and Biogeography 10(3): 305–313. doi – 10.2788/8891

PECBMS (2019). Pan-European Common Bird Monitoring Scheme. Retrieved in 2020-04-10, from https://pecbms.info/.

Pike, N. (2011). Using false discovery rates for multiple comparisons in ecology and evolution. Methods in Ecology and Evolution 2(3): 278–282. doi – 10.1111/j.2041-210X.2010.00061.x

Plieninger, T., Draux, H., Fagerholm, N., Bieling, C., Bürgi, M., Kizos, T., Kuemmerle, T., Primdahl, J. and Verburg, P.H. (2016). The driving forces of landscape change in Europe: A systematic review of the evidence. Land Use Policy 57: 204–214. doi – 10.1016/j.landusepol.2016.04.040

Profico (2008). *Aproveitamento Hidroelétrico de Foz-Tua: Estudo de Impacte Ambiental (EIA)* (Foz-Tua Dam: Environmental Impact Assessment (EIA)). Relatório Técnico, 2 vols. Profico-Ambiente, Lisboa. 357 + 440 pp.

PORDATA (2018). *População residente segundo os Censos: total e por sexo*. (Resident Population according to the Census: total and by sex). PORDATA – Estatísticas, gráficos e indicadores de Municípios, Portugal e Europa. Fundação Francisco Manuel dos Santos, Porto. Retrieved 2020-04-10 2019, from http://www.pordata.pt

R Core Team (2019). R: A Language and Environment for Statistical Computing. R Foundation Vienna, Austria. URL http://www.R-project.org/.

Reif, J. and Vermouzek, Z. (2018). Collapse of farmland bird populations in an Eastern European country following its EU accession. Conservation Letters: e12585. doi – 10.1111/conl.12585

Reif, J., Vorísek, P., Astný, K.S, Bejcek, V. and Petr, J. (2008). Agricultural intensification and farmland birds: new insights from a central European country. Ibis 150(3): 596–605. doi – 10.1111/j.1474-919X.2008.00829.x

Reino, L., Porto, M., Morgado, R., Moreira, F., Fabião, A., Santana, J., Delgado, A., Gordinho, L., Cal, J. and Beja, P. (2010). Effects of changed grazing regimes and habitat fragmentation on Mediterranean grassland birds. Agriculture, Ecosystems & Environment 138(1-2): 27–34. doi – 10.1016/j.agee.2010.03.013

Reino, L., Triviño, M., Beja, P., Araújo, M.B., Figueira, R., & Segurado, P. (2018). Modelling landscape constraints on farmland bird species range shifts under climate change. Science of the Total Environment 625: 1596–1605. doi – 10.1016/j.scitotenv.2018.01.007

Regos A, Domínguez J., Gil-Tena A., Brotons, L., Ninyerola, M. and Pons, X. (2016). Rural abandoned landscapes and bird assemblages: Winners and losers in the rewilding of a marginal mountain area (nw spain) Regional Environmental Change 16: 199–211. doi – 10.1007/s10113-014-0740-7

Ribeiro, J.A. (2000). Caracterização Genérica da Região Vinhateira do Alto Douro (General Characterization of the Upper Douro Wine Region). DOURO–Estudos & Documentos, 5(10): 11–29.

Ribeiro, P.F., Santos, J.L., Bugalho, M.N., Santana, J., Reino, L., Beja, P. and Moreira, F. (2014). Modelling farming system dynamics in High Nature Value Farmland under policy change. *Agriculture*, Ecosystems & Environment 183: 138–144. doi – 10.1016/j.agee.2013.11.002)

Robertson, M.P., Cumming, G.S. and Erasmus, B.F.N. (2010). Getting the most out of atlas data. Diversity and Distributions 16(3): 363–375. doi – 10.1111/j.1472-4642.2010.00639.x

Russo, D. (2006). Effects on Land Abandonment in Europe: Conservation and Management Implications. Università degli Studi di Napoli Federico II, Napoli. .

Sanderson, F.J., Kucharz, M., Jobda, M. and Donald, P.F. (2013). Impacts of agricultural intensification and abandonment on farmland birds in Poland following EU accession. Agriculture, Ecosystems & Environment 168: 16–24. doi – 10.1016/j.agee.2013.01.015

Santana, J., Reino, L., Stoate, C., Borralho, R., Carvalho, C.R., Schindler, S., Moreira, F., Bugalho, M.N., Ribeiro, P.F., Santos, J.L. and Vaz, A. (2014). Mixed effects of long-term conservation investment in Natura 2000 farmland. Conservation Letters 7(5): 467–477. doi – 10.1111/conl.1207

Santana, J., Reino, L., Stoate, C., Moreira, F., Ribeiro, P.F., Santos, J.L., Rotenberry, J.T. and Beja, P. (2017). Combined effects of landscape composition and heterogeneity on farmland avian diversity. Ecology and Evolution 7(4): 1212–1223. doi – 10.1002/ece3.2693

Siegel, S. and Castellan Jr., N.J. (1988). Nonparametric Statistics for the Behavioral Sciences (2nd ed.), McGraw-Hill, New York.

Silva J.P., Correia R., Alonso H., Martins R.C., D’Amico M., Delgado A., Sampaio H., Godinho C., Moreira F. (2018). EU protected area network did not prevent a country wide population decline in a threatened grassland bird. PeerJ 6: e4284. doi – 10.7717/peerj.4284

Sirami, C., Brotons, L. and Martin, J.L. (2007). Vegetation and songbird response to land abandonment: from landscape to census plot. Diversity and Distributions 13(1): 42–52. doi – 10.1111/j.1472-4642.2006.00297.x

Sirami, C., Brotons, L., Burfield, I., Fonderflick, J. and Martin, J.L. (2008). Is land abandonment having an impact on biodiversity? A meta-analytical approach to bird distribution changes in the north-western Mediterranean. Biological Conservation 141(2): 450–459. doi – 10.1016/j.biocon.2007.10.015

Suárez-Seoane, S., Osborne, P.E. and Baudry, J. (2002). Responses of birds of different biogeographic origins and habitat requirements to agricultural land abandonment in northern Spain. Biological Conservation 105(3): 333–344. doi – 10.1016/S0006-3207(01)00213-0

Stoate, C., Boatman, N.D., Borralho, R.J., Carvalho, C.R., De Snoo, G.R. and Eden, P. (2001). Ecological impacts of arable intensification in Europe. Journal of Environmental Management 63(4): 337–365. doi – 10.1006/jema.2001.0473

Stoate, C., Báldi, A., Beja, P., Boatman, N.D., Herzon, I., Van Doorn, A., De Snoo, G.R., Rakosy, L. and Ramwell, C. (2009). Ecological impacts of early 21st century agricultural change in Europe:: a review. Journal of Environmental Management 91(1): 22–46. doi – 10.1016/j.jenvman.2009.07.005

Tibshirani, R. (1996). Regression shrinkage and selection via the lasso. Journal of the Royal Statistical Society. Series B (Methodological): 267–288.

Vorisek, P., Jiguet, F., van Strien, A., Skorpilova, J., Klvanova, A. and Gregory, R.D. (2010). Trends in abundance and biomass of widespread European farmland birds: how much have we lost. BOU *Proceedings*, Lowland Farmland Birds III: 1–24.

Wang, Y.I., Naumann, U., Wright, S.T. and Warton, D.I. (2012). mvabund–an R package for model-based analysis of multivariate abundance data. Methods in Ecology and Evolution 3(3): 471–474. doi – 10.1111/j.2041-210X.2012.00190.x

Wretenberg, J., Lindström, Å., Svensson, S. and Pärt, T. (2007). Linking agricultural policies to population trends of Swedish farmland birds in different agricultural regions. Journal of Applied Ecology 44(5): 933–941. doi – 10.1111/j.1365-2664.2007.01349.x

Zakkak, S., Radovic, A., Nikolov, S.C., Shumka, S., Kakalis, L. and Kati, V. (2015). Assessing the effect of agricultural land abandonment on bird communities in southern-eastern Europe. Journal of Environmental Management 164: 171–179. doi – 10.1016/j.jenvman.2015.09.005

