## Supplementary material for "Mind the gap: assessing the co-occurrying impacts of intensification and land abandonment on bird diversity in a farmland mosaic of Northeastern Portugal using historical data (1980–2000)": Includes Supplementary Figures S1-S3 and Supplementary Tables ST1 and ST2

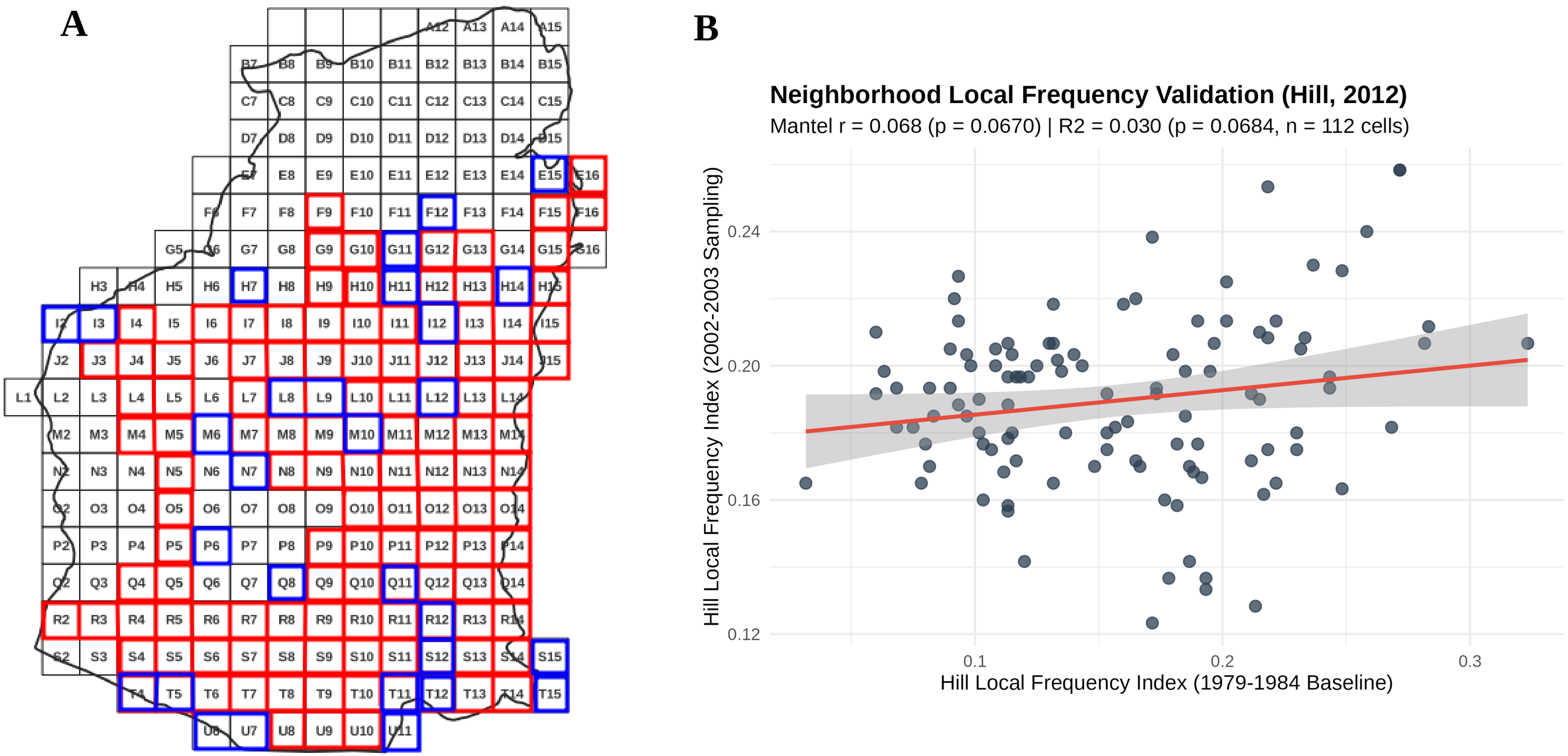


**Figure S1.** Assessment of spatial sampling coverage and Neighborhood Local Frerquency Validation (Hill, 2012) between atlas campaigns. (A) Geographical distribution of the n=112 paired 500m×500m grid squares sampled in both Period 1 (1979–1984) and Period 2 (2002–2003) (red-bordered squares), or only in Period 1 (blue-bordered squares). All squares, including those not sampled (black-bordered squares) were used for the landscape global analysis. (B) Neighborhood-weighted spatial validation following Hill's (2012) framework. Relationship between the Neighborhood Local Frequency Index in Period 1 (x-axis) and Period 2 (y-axis) across n=112 overlapping cells for 75 shared species (k=8 nearest neighbors). The red line represents the linear regression fit (R2=0.030, p=0.0680). The spatial validation (Mantel r=0.067, p=0.0680) confirms that temporal dynamics represent genuine ecological turnover rather than an artifact of reduced sampling duration in Period 2.

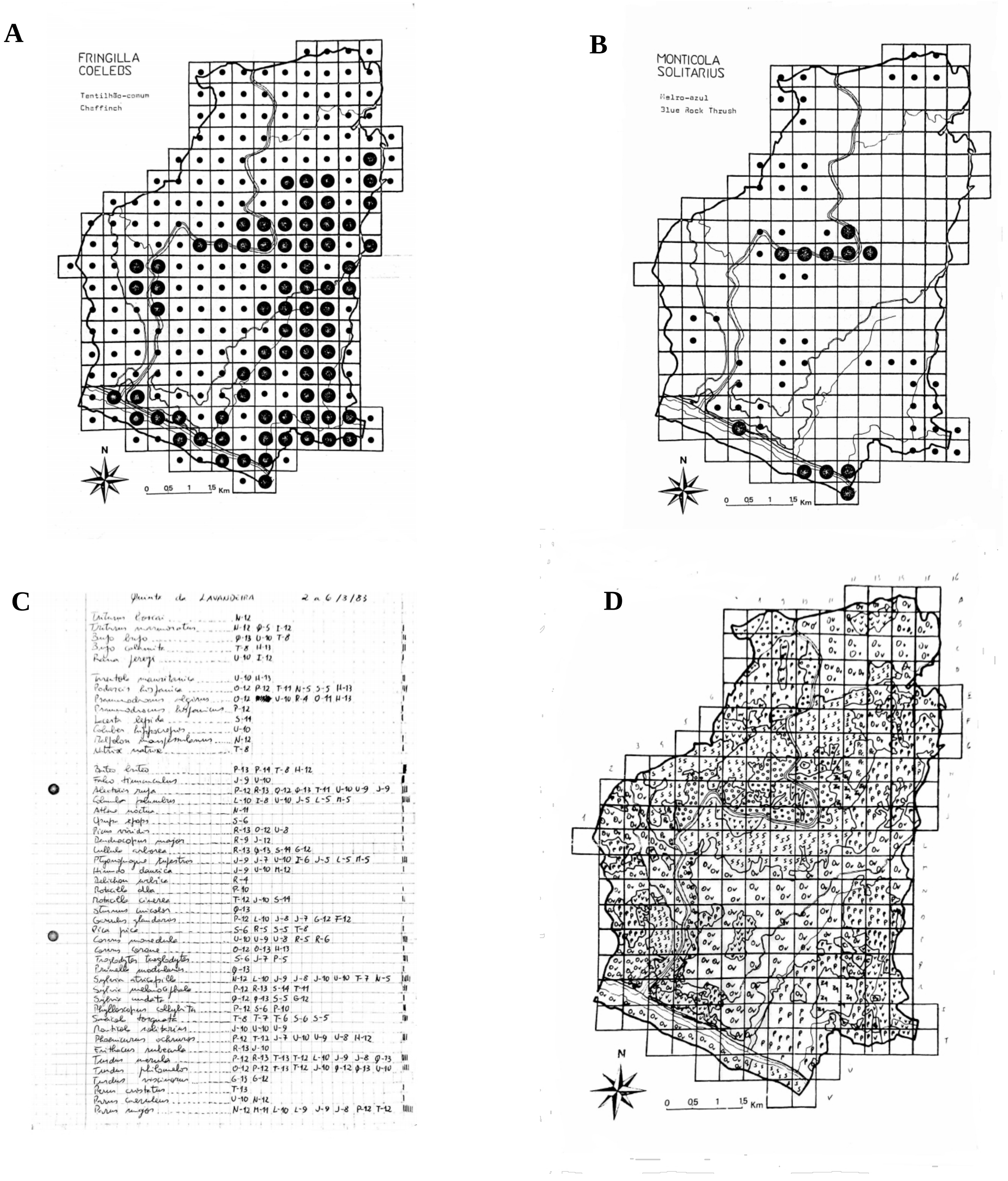


**Figure S2.** Examples of the data information in Ferrand de Almeida (unpublished) manuscript: species maps for **(A)** an abundant species (Fringilla coelebs, and for **(B)** a species with a more restricted abundance (Monticola solitarius); **(C)** a raw datasheet (the only surviving), with the layout of the Vertebrate species for a sampling occasion, with the identification of the squares were they were observed, and **(D)** the original map with the landscape characteritsics of each square in 1980 (the letters denote the main land-use classes used in the paper, with some adaptations). In the species maps, the large dots represent actual presence, while the small dots represents squares of potential presence, according to the species habitat requirements (see Table 2 in Supplementary Material) and the occurrence of those habitats in each square.

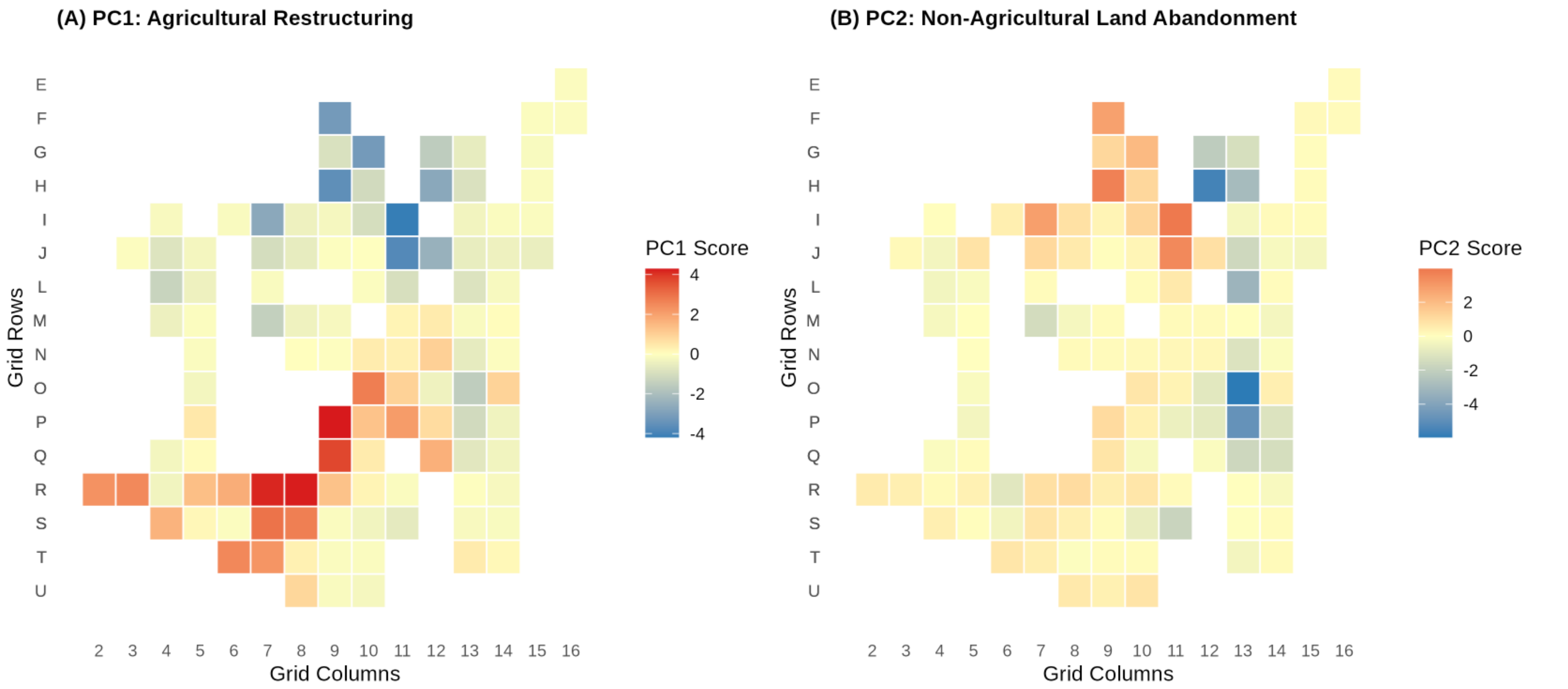


**Figure S3**. Spatial distribution of Principal Component Analysis (PCA) scores across the 500m×500m sampling grid squares (n=112), summarizing landscape dynamics between 1980 and 2000. (A) PC1 axis: (Agricultural Restructuring): Positive values (red) indicate areas dominated by agricultural restructuring and intensive vineyard expansion, whereas negative values (blue) represent agricultural abandonment (loss of traditional mixed vineyards and olive orchards. (B) PC2 axis (Non-Agricultural Land Abandonment & Natural Succession): Positive values (red) depict semi-natural abandoned areas characterized by shrubland expansion, either natural or as a consquence of fire (burnt areas) and vegetation recovery (concentrated in the northern grid cells), whereas negative values (blue) represent pinewoods and areas with lower succession activity. The square grid labels correspond to the spatial sampling layout (Rows E–U; Columns 2–16), rather than to the entire study area.

**Table S1.** List of all species observed in the study area in 1980-84 and in 2002-03, classified by **biogeographic origin** (E='Eurosiberian', M='Mediterranean'), **phenology** (R='Resident', W='Winter visitort', S='Spring visitor', Ac ='Accidental') and **habitat** (Dc='Dc', Sc='Sc', DC='Dense shrublands', O='Open habitats'; Other habitats: Wb='Water bodies', RW='RW', R='Natural R', R/U='Natural rocky and urban areas', Ec='Ec' ). **Abundace in 1980**: class of abundance as in Ferrand de Almeida (unpublished) ('Rare': 1-20 birds/couples, 'Common': 20-200, 'Abundant': >200). When available, the exact number of couples ('c') or individuals ('i') are given in brackets. The number of 500x500 m squares in which a species was observed in each period is presented (total N=112), the binomial deviance (the parameter dDi), a measure of the uncertainity in the occurrence of a particular species between the two periods, and its level of significance, adjusted by the False discovery Rate (FDR) method (statistical significant values in bold). A dDi<0 indicates a "community-convergence" species; a dDi>0, a "communtiy-divergence" species (see text for details). ni = not included. The species of sporadic occurrence are marked with a (+). Only the species marked with a (*), were used in the 4th-corner analysis.

| ***Families/Species*** | ***Code*** | ***Common Names*** | ***Origin*** | ***Phen.*** | ***Habitat***[2] | ***Abundance in 1980***[2] | ***N. of squares*** | | ΔDi | ***p*** |
|---|---|---|---|---|---|---|---|---|---|---|
| | | | | | | | 1979-84 | 2002-03 | | |
| **Fam. Phalacrocoraxidae** | | | | | | | | | | |
| *Phalacrocorax carbo* + | *Pcar* | Cormorant | E | W | Wb | - | 0 | 2 | +20.07 | *0.67* |
| **Fam. Ardeidae** | | | | | | | | | | |
| *Ardea cinerea* + | *Acin* | Grey Heron | E | W | Wb | Common | 24 | 3 | -88.75 | ***0.03*** |
| **Fam. Accipitridae** | | | | | | | | | | |
| *Neophron percnopterus* + | *Nper* | Egyptian Vulture | M | S | R | Rare (1c) | 3 | 0 | -27.64 | *0.49* |
| *Aquila chrysaetos*+ | *Achr* | Golden Eagle | E | Sp | R | Rare | 1 | 1 | 0.00 | *1.00* |
| *Hieraaetus fasciatus* + | *Hfas* | Bonelli's Eagle | M | R | R | Rare (1c) | 15 | 2 | -68.14 | ***0.03*** |
| *Buteo buteo* | *Bbut* | Buzzard | E | R | Sc | Rare (5-6c) | 32 | 6 | -87.22 | ***0.03*** |
| *Accipiter nisus* | *Anis* | Sparrowhawk | E | W | Dc | Rare | 5 | 4 | -6.35 | *1.00* |
| *Accipiter gentilis* | *Agen* | Goshawk | E | R | Dc | Rare (2c) | 8 | 2 | -35.57 | *0.12* |
| *Milvus migrans* + | *Mmig* | Black Kite | E | Sp | Sc | Rare | 2 | 1 | 0.00 | *1.00* |
| *Circus pygargus* + | *Cpyg* | Montagu's Harrier | E | Sp | O | - | 0 | 1 | 0.00 | *1.00* |

| *Families/Species* | *Code* | *Common Names* | *Origin* | *Phen.* | *Habitat*[2] | *Abundance in 1980*[2] | *N. of squares* | | ΔDi | *p* |
|---|---|---|---|---|---|---|---|---|---|---|
| *Circaetus gallicus* + | *Cgal* | Short-toed Eagle | M | Sp | Sc | - | 0 | 1 | +11.43 | *1.00* |
| *Falco tinnunculus* | *Ftin* | Kestrel | E | R | Sc | Rare | 16 | 12 | -15.59 | *0.44* |
| *Falco columbarius* + | *Fcol* | Merlin | E | Sp | Sc | Rare | 2 | 0 | +20.07 | *0.73* |
| *Falco naumanii* + | *Fnau* | Lesser Kestrel | E | Sp | Sc | Rare | 2 | 0 | +20.07 | *0.73* |
| **Fam. Phasianidae** | | | | | | | | | | |
| *Alectoris rufa* * | *Aruf* | Red-legged Partridge | M | R | O | Abundant | 30 | 5 | -89.31 | ***0.03*** |
| **Fam. Laridae** | | | | | | | | | | |
| *Larus fuscus* | *Lfus* | Lesser Black-backed Gull | E | W | Wb | Rare | 1 | 0 | -11.43 | *1.00* |
| *Larus ridibundus* | *Lrid* | Black-headed Gull | E | W | Wb | Abundant | 10 | 0 | -67.40 | ***0.03*** |
| **Fam. Charadriidae** | | | | | | | | | | |
| *Actitis hypoleucos* + | *Ahyp* | Common Sandpiper | E | Sp | Wb | Rare | 1 | 0 | -11.43 | *1.00* |
| **Fam. Columbidae** | | | | | | | | | | |
| *Columba palumbus* * | *Cpal* | Woodpigeon | E | R | Dc | Abundant. | 52 | 33 | -17.09 | *0.03* |
| *Columba livia* + | *Cliv* | Rock dove | E | Sp | Dc | Rare | 1 | 0 | +16.21 | 1.00 |
| *Streptopelia turtur* * | *Stur* | Turtle Dove | E | S | Sc | Common | 14 | 34 | +53.11 | ***0.03*** |
| **Fam. Cuculidae** | | | | | | | | | | |
| *Cuculus canorus* * | *Ccan* | Cuckoo | E | S | Sc | Common | 12 | 6 | -29.48 | *0.43* |
| **Fam. Strigidae** | | | | | | | | | | |
| *Otus scops* | *Osco* | Scops Owl | M | S | Sc | Common | 6 | 1 | -26.73 | *0.31* |
| *Athene noctua* | *Anoc* | Little Owl | M | R | Sc | Rare | 12 | 2 | -56.21 | ***0.03*** |
| *Asio otus* + | *Aotu* | Long-eared Owl | E | Sp | Dc | Rare | 1 | 0 | -11.43 | *1.00* |
| **Fam. Caprimulgidae** | | | | | | | | | | |
| *Caprimulgus europaeus* | *Ceur* | Nightjar | E | S | Ec | Rare | 5 | 1 | -29.44 | *0.31* |

| *Families/Species* | *Code* | *Common Names* | *Origin* | *Phen.* | *Habitat*[2] | *Abundance in 1980*[2] | *N. of squares* | | ΔDi | *p* |
|---|---|---|---|---|---|---|---|---|---|---|
| **Fam. Apodidae** | | | | | | | | | | |
| *Apus apus* + | *Aapu* | Swift | E | Sp | R/U | - | 0 | 1 | +11.43 | *1.00* |
| **Fam. Alcedinidae** | | | | | | | | | | |
| *Alcedo atthis* | *Aatt* | Kingfisher | E | R | Wb | Common | 5 | 4 | -6.35 | *1.00* |
| **Fam. Meropidae** | | | | | | | | | | |
| *Merops apiaster* * | *Mapi* | Bee-eater | M | S | RW | Rare | 4 | 5 | +6.35 | *1.00* |
| **Fam. Upupidae** | | | | | | | | | | |
| *Upupa epops* * | *Uepo* | Hoopoe | E | S | Sc | Common | 13 | 12 | -4.15 | *1.00* |
| **Fam. Picidae** | | | | | | | | | | |
| *Picus viridis* * | *Pvir* | Green Woodpecker | E | R | Dc | Common | 35 | 16 | -47.26 | ***0.03*** |
| *Dendrocopus major** | *Dmaj* | Great Spotted Woodpecker | E | R | Dc | Common | 22 | 29 | +19.20 | *0.43* |
| *Jynx torquilla* + | *Jtor* | Wryneck | E | S | Ec | Rare | 0 | 1 | +20.07 | 0.73 |
| **Fam. Alaudidae** | | | | | | | | | | |
| *Lululla arborea* * | *Larb* | Woodlark | E | W | O | Common | 6 | 13 | +33.63 | *0.26* |
| *Alauda arvensis* + | *Aarv* | Skylark | E | Sp | O | Rare | 3 | 0 | -27.64 | 0.54 |
| **Fam. Hirundinidae** | | | | | | | | | | |
| *Ptyonoprogne rupestris* | *Prup* | Sand Martin | M | R | R | Common | 40 | 27 | -20.03 | *0.07* |
| *Hirundo rustica* | *Hrus* | Swallow | E | S | R/U | Common | 10 | 20 | +37.71 | *0.17* |
| *Hirundo daurica* | *Hdau* | Red-rumped Swallow | E | S | R | Common | 14 | 22 | +26.58 | *0.32* |
| *Delichon urbica* | *Durb* | House Martin | E | S | R/U | Abundant | 22 | 19 | -8.98 | *0.91* |
| **Fam. Motacilidae** | | | | | | | | | | |
| *Anthus pratensis* * | *Apra* | Meadow Pipit | E | W | O | Abundant | 6 | 5 | -5.93 | *1.00* |
| *Motacilla cinerea* * | *Mcin* | Grey Wagtail | E | R | RW | Common | 15 | 9 | -11.94 | *0.87* |

| *Families/Species* | *Code* | *Common Names* | *Origin* | *Phen.* | *Habitat*[2] | *Abundance in 1980*[2] | *N. of squares* | | ΔDi | *p* |
|---|---|---|---|---|---|---|---|---|---|---|
| *Motacilla alba* * | *Malb* | White Wagtail | E | W | O | Common | 13 | 29 | +53.59 | ***0.03*** |
| **Fam. Lanidae** | | | | | | | | | | |
| *Lanius meridionalis* * | *Lmer* | Great Grey Shrike | E | R | O | Common | 8 | 8 | 0.00 | *1.00* |
| *Lanius senator* + | *Lsen* | Woodchat Shrike | M | S | O | Rare | 2 | 1 | -8.64 | *1.00* |
| **Fam. Oriolidae** | | | | | | | | | | |
| *Oriolus oriolus* * | *Oori* | Golden Oriole | E | S | Sc | Common | 25 | 19 | -16.94 | *0.56* |
| **Fam. Sturnidae** | | | | | | | | | | |
| *Strunus unicolor* * | *Suni* | Spotless Starling | M | R | O | Common | 8 | 18 | +41.11 | *0.08* |
| *Sturnus vulgaris* + | *Svul* | Starlin | E | Sp | O | Rare | 1 | 0 | -11.43 | 1.00 |
| **Fam. Corvidae** | | | | | | | | | | |
| *Garrulus glandarius* * | *Ggla* | Jay | E | R | Sc | Abundant | 47 | 22 | -28.64 | ***0.05*** |
| *Pica pica* * | *Ppic* | Magpie | E | R | Sc | Common | 16 | 5 | -51.00 | ***0.03*** |
| *Corvus corone* * | *Ccor* | Corvus corone | E | R | Sc | Common | 15 | 4 | -53.70 | ***0.05*** |
| *Corvus monedula* | *Cmon* | Jackdaw | E | R | Sc | Abundant | 13 | 0 | -80.42 | *0.03* |
| *Corvus corax* | *Ccorx* | Raven | E | R | Dc | Rare | 4 | 1 | -23.09 | *0.41* |
| **Fam. Cinclidae** | | | | | | | | | | |
| *Cinclus cinclus* | *Ccin* | Dipper | E | R | Wb | Rare | 6 | 1 | -35.37 | *0.16* |
| **Fam. Troglodytidae** | | | | | | | | | | |
| *Troglodytes troglodytes* * | *Ttro* | Wren | E | R | DC | Abundant | 11 | 44 | +80.42 | ***0.03*** |
| **Fam. Prunellidae** | | | | | | | | | | |
| *Prunella modularis** | *Pmod* | Dunnock | E | W | DC | Abundant | 12 | 3 | -48.63 | ***0.05*** |
| **Fam. Sylvidae** | | | | | | | | | | |
| *Cettia cetti* * | *Ccet* | Cetti's Warbler | M | R | RW | Common | 4 | 12 | +41.76 | *0.08* |

| ***Families/Species*** | ***Code*** | ***Common Names*** | ***Origin*** | ***Phen.*** | ***Habitat[2]*** | ***Abundance in 1980[2]*** | ***N. of squares*** | | ΔDi | ***p*** |
|---|---|---|---|---|---|---|---|---|---|---|
| *Cisticola juncidis* + | *Cjun* | Fan-tailed Warbler | E | Sp | RW | Rare | 1 | 0 | -11.43 | 1.00 |
| *Sylvia atricapilla* * | *Satr* | Blackcap | E | R | DC | Abundant | 31 | 63 | +17.95 | ***0.03*** |
| *Sylvia melanocephala* * | *Smel* | Sardinian Warbler | M | R | DC | Abundant | 25 | 85 | -7.96 | ***0.03*** |
| *Sylvia undata* * | *Sund* | Dartford Warbler | M | R | DC | Abundant | 23 | 8 | -56.09 | ***0.03*** |
| *Sylvia cantillans* + | *Scan* | Subalpine Warbler | M | S | DC | Rare | 4 | 3 | -6.88 | 1.00 |
| *Sylvia communis* + | *Scom* | Sylvia communis | E | Sp | DC | Rare | 3 | 0 | -27.64 | 0.54 |
| *Hippolais polyglotta* | *Hpol* | Melodious Warbler | M | S | O | Rare | 3 | 2 | -7.57 | *1.00* |
| *Phylloscopus collybita* * | *Pcol* | Chiffchaff | E | W | DC | Abundant | 26 | 16 | -29.51 | *0.17* |
| *Regulus ignicapillus* * | *Rign* | Firecrest | E | R | Dc | Rare | 1 | 14 | +72.97 | *0.03* |
| **Fam. Turdidae** | | | | | | | | | | |
| *Oenanthe hispanica* + | *Ohis* | Black-eared Wheatear | M | S | O | Rare | 1 | 1 | 0.00 | *1.00* |
| *Oenanthe leucura* * | *Oleu* | Black Wheatear | M | R | O | Rare | 1 | 10 | +46.21 | 0.08 |
| *Saxicola torquata* * | *Stor* | Stonechat | E | R | O | Abundant | 22 | 25 | +7.96 | *0.94* |
| *Monticola solitarius* | *Msol* | Blue Rock Trush | M | R | R | Common | 10 | 18 | +34.59 | *0.16* |
| *Phoenicuros ochruros* * | *Poch* | Black Redstart | M | R | R/U | Common | 30 | 54 | +24.95 | ***0.03*** |
| *Erithacus rubecula* * | *Erub* | Robin | E | W/R[1] | Sc | Abundant | 25 | 98 | -42.66 | ***0.03*** |
| *Luscinia megarhynchos* * | *Lmeg* | Nightingale | E | S | RW | Common | 10 | 14 | +17.00 | *0.61* |
| *Turdus merula* * | *Tmer* | Blackbird | E | R | Sc | Abundant | 70 | 101 | -90.55 | ***0.03*** |
| *Turdus philomelos* * | *Tphi* | Song Thrush | E | W | Sc | Abundant | 49 | 26 | -32.14 | ***0.03*** |
| *Turdus illiacus* + | *Till* | Redwing | E | W | Sc | Common | 1 | 2 | +8.64 | *1.00* |
| *Turdus viscivorus* * | *Tvis* | Mistle Thrush | E | R | Sc | Common | 9 | 20 | +42.27 | *0.07* |
| *Turdus pilaris* + | *Tpil* | Fieldfire | E | W | Sc | Common | 1 | 1 | 0.00 | *1.00* |

| *Families/Species* | *Code* | *Common Names* | *Origin* | *Phen.* | *Habitat*[2] | *Abundance in 1980*[2] | *N. of squares* | | ΔDi | *p* |
|---|---|---|---|---|---|---|---|---|---|---|
| **Fam. Paridae** | | | | | | | | | | |
| *Parus cristatus* * | *Pcri* | Crested Tit | E | R | Dc | Common | 3 | 30 | +106.37 | ***0.03*** |
| *Parus caeruleus* * | *Pcae* | Blue Tit | E | R | Dc | Abundant | 19 | 52 | +52.70 | ***0.03*** |
| *Parus ater* * | *Pate* | Coal Tit | E | R | Dc | Abundant | 5 | 26 | +82.85 | ***0.03*** |
| *Parus major* * | *Pmaj* | Great Tit | E | R | Dc | Abundant | 49 | 72 | -8.73 | ***0.05*** |
| *Aeghitalos caudatus* * | *Acau* | Long-tailed Tit | E | R | Dc | Abundant | 25 | 28 | +7.03 | *0.99* |
| **Fam. Sittidae** | | | | | | | | | | |
| *Sitta europaea* * | *Seur* | Nuthatch | E | R | Dc | Common | 1 | 12 | +64.84 | ***0.03*** |
| **Fam. Certhidae** | | | | | | | | | | |
| *Certhia brachydactyla* * | *Cbra* | Short-toed Treecreeper | E | R | Dc | Abundant | 7 | 49 | +101.14 | ***0.03*** |
| **Fam. Passeridae** | | | | | | | | | | |
| *Passer domesticus* * | *Pdom* | House Sparrow | E | R | R/U | Abundant | 22 | 25 | +7.96 | *0.74* |
| Passer montanus * | *Pmon* | Tree Sparrow | E | R | O | Common | 3 | 7 | +24.76 | 0.56 |
| *Petronia petronia* + | *Ppet* | Rock Sparrow | M | S | O | Rare | 2 | 1 | -8.64 | *1.00* |
| **Fam. Fringillidae** | | | | | | | | | | |
| *Fringilla coelebs* * | *Fcoe* | Chaffinch | E | R | Sc | Abundant | 67 | 90 | -39.94 | ***0.03*** |
| *Pyrrhula pyrrhula* | *Ppyr* | Bullfinch | E | W | Sc | Common | 3 | - | -27.64 | *0.43* |
| *Serinus serinus* * | *Sser* | Serin | M | R | Sc | Abundant | 50 | 79 | -18.18 | ***0.03*** |
| *Carduelis chloris* * | *Cchl* | Greenfinch | E | R | Sc | Abundant | 12 | 49 | +77.70 | ***0.03*** |
| *Carduelis spinus* * | *Cspi* | Siskin | E | W | Sc | Abundant | 5 | 7 | +11.51 | *0.91* |
| *Carduelis carduelis* * | *Ccar* | Goldfinch | E | R | Sc | Abundant | 17 | 59 | +58.13 | ***0.03*** |
| *Carduelis cannabina* * | *Ccann* | Linnet | E | R | Sc | Abundant | 5 | 28 | +85.10 | ***0.03*** |
| *Miliaria calandra* + | *Mcal* | Corn Bunting | E | R | O | Common | 1 | 1 | 0..00 | *1.00* |

| ***Families/Species*** | ***Code*** | ***Common Names*** | ***Origin*** | ***Phen.*** | ***Habitat***[2] | ***Abundance in 1980***[2] | ***N. of squares*** | | ΔDi | ***p*** |
|---|---|---|---|---|---|---|---|---|---|---|
| *Coccothraustes coccothraustes* * | *Ccoc* | Hawfinch | E | Sp | Dc | - | 0 | 4 | +34.51 | 0.21 |
| **Fam. Emberizidae** | | | | | | | | | | |
| *Emberiza cia* * | *Ecia* | Rock Bunting | E | R | R/U | Abundant | 43 | 46 | +5.20 | *0.46* |
| *Emberiza cirlus* * | *Ecir* | Cirl Bunting | M | R | O | Abundant | 17 | 15 | -7.17 | *1.00* |
| **Fam. Estrildidae** | | | | | | | | | | |
| *Estrilda astrild* * | *East* | Waxbill | Ex | R | RW | - | 0 | 7 | +62.64 | 0.06 |

[1]This species was a winter visitor in 1980 (Ferrand de Almeida, unpublished), but turned into a resident species (see text)
[2]After Ferrand de Almeida (unpublished)

**Table S2.** List and description the ecological parameters used in the study to characterize the ecological characteristics of the global bird assemblages (Table 2). The **habitat** categories are in accordance to Ferrand de Almeida (unpublished). In brackets are the pooling categories used in the text. The **biogeographic origin** is adapted from Suárez-Seoane et al. (2002); **Phenology** status in Portugal, from Catry et al. (2010) ) and **diet** from Cramp & Simmons (1998) and Catry et al. (2010) for the general patterns and Ferrand de Almeida et al (unpublished) for some specific local cases.

| Parameter | Code | Description |
|---|---|---|
| **Habitat type** | | |
| Sparse tree cover (farmland) | Sc | Traditional agricultural habitats with sparse trees (olive-orchards, vineyards) and degraded natural habitats (e.g., immediate post-fire areas with low amounts of scrubland) |
| Dense tree cover (woodland) | Dc | Large and continuous stretches of woodlands of all types |
| Open areas | O | Open agricultural habitats (e.g., intensive vineyards, orchards and gardens) and open natural habitats (e.g., wet meadow) |
| Dense shrublands | DS | Mainly *Cistus* shrublands, either arbustive or semi-arboreal, but other types as well (e.g., *Genista* spp.) |
| Rocky areas (other) | R | Mainly natural rocky habitats, with dense (but low) shrublands and/or some trees |
| Rocky/urban areas (other) | R/U | Mainly bare rocky habitats or human-made rocky habitats (e.g., vineyard walls, stone building, abandoned or not, or warehouses |
| Riverine woodlands (other) | RW | Woodland habitats in the margins of waterlines |
| Ecotones (other) | Ec | Transition zones between agricultural and woodland areas |
| Water bodies (other) | Wb | River beds, streams, ponds |
| **Biogeographic origin** | | |
| Eurosiberian | E | Large-scale distribution in the Holartic (ie, Northern hemisphere) or Paleartic regions |
| Mediterranean | M | Mediterranean distribution, *sensu latu*, ie., circum-Mediterranean distribution and associated steppic or pseudo-steppe regions with the same climatic characteristics (Middle East). |
| Alloctonous | A | Distribution outside the two previous areas (eg., introduced African species). |

| | | |
|---|---|---|
| **Phenology** | | |
| Resident | R | Present all year round. |
| Winter visitor | W | Present in the winter months (November-March). |
| Summer visitor | S | Present in the summer months (April-July). |
| Accidental | A | Sporadic occurrence in the study area. |
| **Diet** | | |
| Granivorous | G | Mainly grains and berries |
| Insectivorous | I | Mainly insects |
| Mixed diet | M | Mainly insects, but shifting to other food sources in some parts of the year. |
| Omnivorous | O | All sources of food, depending on their availability. |
